# SHANK3-actin cytoskeleton coupling regulates nanoscale remodelling of the postsynaptic density

**DOI:** 10.64898/2026.11.26.769461

**Authors:** Wouter J. Droogers, Albert K. Serweta, Lieke C.P. Steijvers, Amy A. Moerkerken, Oscar K. Klock, Florian Berger, Harold D. MacGillavry

## Abstract

Dynamic remodelling of multiprotein signalling complexes allows cells to rapidly adapt signalling strength. At synapses, neurotransmitter receptors and scaffolds concentrate in subsynaptic nanoclusters that govern synaptic signalling strength and that are dynamically remodelled by synaptic activity. However, although this distinct nanoscale organization of synaptic protein complexes is well-established, the mechanisms that control the activity-induced reorganization of these complexes remain elusive. Here, to resolve the mechanisms underlying the activity-induced spatiotemporal reorganization of the PSD, we combined CRISPR/Cas9-mediated tagging of endogenous synaptic proteins and quantitative super-resolution microscopy. We found that synaptic potentiation drives a rapid and long-lasting increase in the number and density of PSD scaffold nanoclusters. Strikingly, activity-induced reorganization of scaffold proteins originated in deep PSD layers and preceded AMPA receptor repositioning, indicating that newly recruited receptors populate preformed scaffold nanoclusters. Computational simulations support that this reorganization is sufficient to drive potentiation of synaptic responses. Mechanistically, we found that the interaction between the scaffold SHANK3 and the actin cytoskeleton is essential for activity-induced PSD nanocluster remodelling. Altogether, our findings present a novel mechanism through which cytoskeletal interactions remodel the spatial organization of the PSD, advancing our understanding of how the dynamic nanoscale reorganization of protein complexes can drive neuronal signalling.

**Highlights:** • Synaptic activity induces nanoscale redistribution of synaptic protein complexes

• Scaffold reorganization precedes AMPA receptor repositioning

• Increasing nanocluster density can facilitate synaptic potentiation

• SHANK3-actin interactions drive activity-induced nanocluster reorganization

## Introduction

Synaptic strength is critically determined by the composition and nanoscale organization of multi-protein assemblies in the postsynaptic density (PSD). At excitatory synapses, AMPA receptors and multi-domain scaffold molecules concentrate in subsynaptic nanoclusters,^1–3^ that preferentially align with presynaptic release sites forming transsynaptic ‘nanocolumns’.^4^ Importantly, this organization critically controls synaptic strength by maximizing receptor activation in response to glutamate release.^5,6^ Alterations in synapse nanostructure could thus support long-term changes in synaptic strength.^7^ Indeed, induction of long-term potentiation (LTP) was found to alter PSD nano-organization.^4,8^ However, how this refined level of subsynaptic reorganization is modulated during plasticity to alter synaptic strength remains elusive.

AMPA receptors are anchored at the PSD via interactions with multidomain scaffolding proteins. Particularly, the prototypical scaffolding protein PSD95 is central in retaining and positioning AMPA receptors at the synaptic membrane.^9^ PSD95, in turn, interacts with GKAP scaffolds deeper in the PSD that couple to SHANK proteins.^10^ The SHANK scaffolding platform forms the deepest layer of the PSD that contacts the actin cytoskeleton directly and via diverse actin regulators.^11–14^ Thus, the actin cytoskeleton can engage at the base of the nanocolumn, and activity-dependent actin dynamics could drive rearrangements of nanostructure within the PSD. However, how the actin cytoskeleton controls plasticity-induced reorganization of the PSD remains unexplored.

Here, we used CRISPR/Cas9-mediated labelling of endogenous synaptic proteins and single-molecule localization microscopy (SMLM),^15,16^ to demonstrate that synaptic activity initiates the reorganization of subsynaptic scaffold nanoclusters in the PSD ultimately leading to repositioning of AMPA receptors. We observed a rapid increase in the number and density of nanoclusters formed by the scaffolds PSD95, GKAP and SHANK3, and GluA1-containing AMPA receptors, even though PSD size did not increase in the early phases of LTP. Computational simulations indicated that high nanocluster density facilitates the potentiation of synaptic responses. Furthermore, using knock-down and rescue experiments we found that SHANK3, and specifically its interaction with the actin cytoskeleton, is required for the activity-induced reorganization of the PSD.

## Results

### Synaptic activity induces the spatiotemporal reorganization of scaffolding nanoclusters within the PSD

The scaffolding proteins PSD95, GKAP and SHANK are among the most abundant structural proteins in the PSD and all form subsynaptic nanoclusters. To test how the distribution of these proteins within the PSD is modulated during LTP, we developed CRISPR/Cas9 knock-in constructs to tag endogenous PSD95, GKAP, SHANK3 and the AMPA-receptor subunit GluA1 with HaloTag allowing for versatile labelling (Figure S1).

To measure temporal changes in the subsynaptic positioning of these key synaptic components, we used direct stochastic optical reconstruction microscopy (dSTORM) before and at different time points after cLTP induction (Figures 1A and 1B). We confirmed that cLTP induction increased surface AMPA receptor levels and dendritic spine growth (Figures S2A-C). Nanoclusters were identified using a local-density-based cluster analysis in each PSD marked by Homer1c (Figure 1B). Interestingly, we found that already 15 min after cLTP induction, the number of PSD95 nanoclusters per PSD was significantly increased (Figures 1C and 1D). This increase was detectable up to 55 min after cLTP induction. Moreover, the density of these nanoclusters, relative to the molecular density in the entire PSD, was also increased (Figure 1E). Similarly, for GKAP we observed an increase in the number of nanoclusters 15 min after cLTP induction (Figures 1F and 1G), which decreased slightly after 55 min. The density of GKAP nanoclusters did not significantly increase (Figure 1H). Remarkably, the number of SHANK3 nanoclusters per PSD was increased immediately after cLTP induction (Figures 1I and 1O). The density of SHANK3 nanoclusters peaked at 15 min but did not persist until 55 min (Figure 1K). In contrast, GluA1-containing AMPA receptors concentrated into more nanoclusters with a higher density only 55 min after cLTP induction (Figures 1L-N). In conclusion, the PSD scaffolding core internally reorganizes quickly after LTP, preceding subsynaptic repositioning of GluA1-containing AMPA receptors.

**Figure 1:**
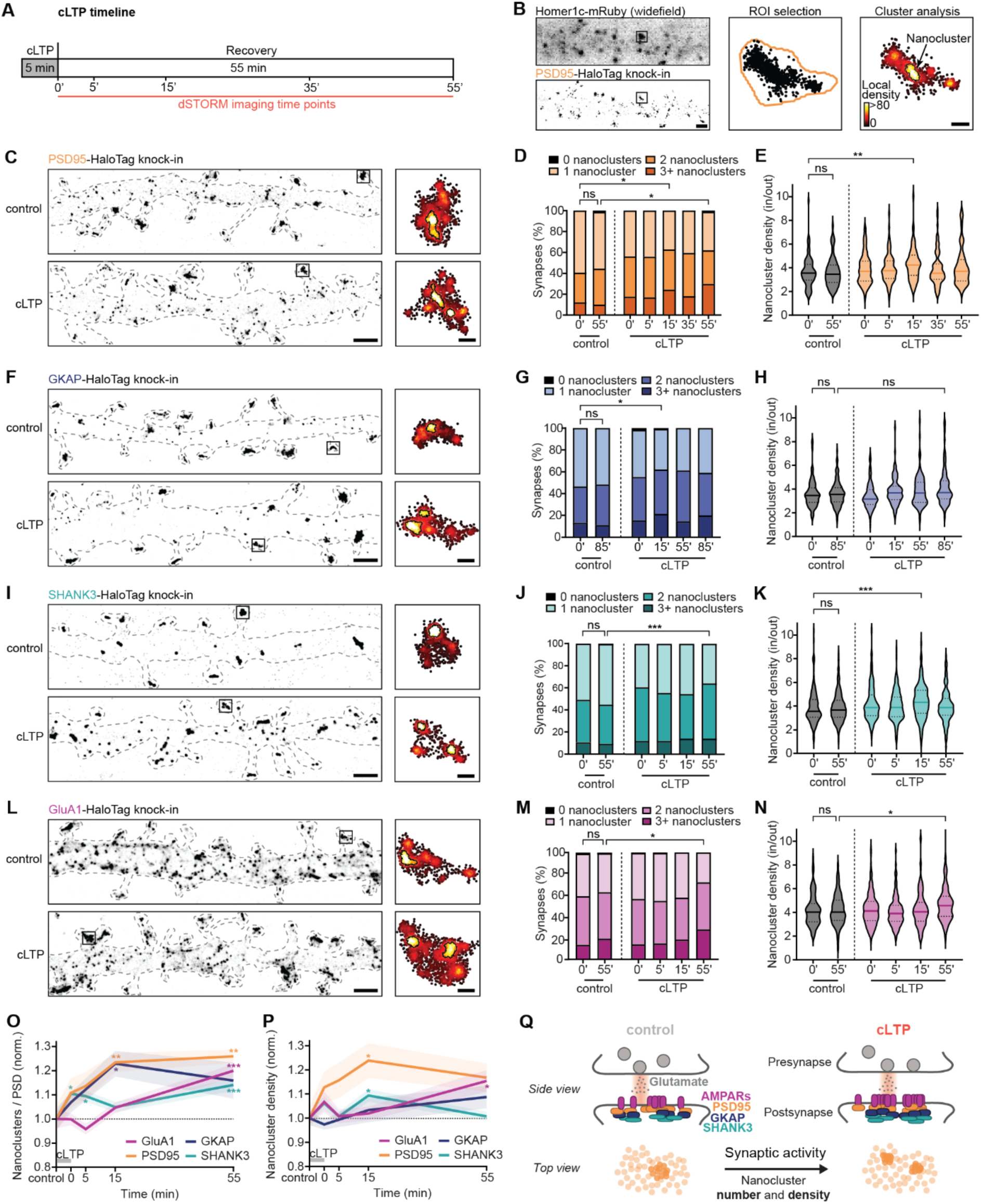
cLTP induces spatiotemporal reorganization of postsynaptic scaffolding and AMPA receptor nanoclusters. **(A)** Timeline of cLTP experiment. **(B)** Left: A representative dendritic stretch of a dissociated hippocampal neuron, visualized with a widefield image of the PSD marker Homer1c and a reconstructed dSTORM image of PSD95-HaloTag. Scale bar: 1 µm. Right: Example of a PSD after nanocluster analysis. Scale bar: 200 nm. **(C)** Left: Representative dendritic stretches 0 and 15 minutes after cLTP induction. Scale bar: 1 µm. Right: PSDs indicated with black squares are shown after nanocluster analysis. Scale bar: 200 nm. **(D)** The number of PSD95 nanoclusters per PSD before and after cLTP induction. N=80-104 synapses from 10-13 neurons. **(E)** Violin plot of the relative density of PSD95 nanoclusters before and after cLTP induction. Sample sizes are the same as in D. **(F)** Same as in C but for GKAP-HaloTag. Examples of control (85 min) and cLTP (15 min). **(G)** Same as in D but for GKAP nanoclusters. N=100-180 synapses from 10-18 neurons. **(H)** Same as in E but for GKAP nanoclusters. Sample sizes are the same as in H. **(I)** Same as in C but for SHANK3-HaloTag. Examples of control (0 min) and cLTP (0 min). **(J)** Same as in D but for SHANK3 nanoclusters. N=156-208 synapses from 11-16 neurons. **(K)** Same as in D but for SHANK3 nanoclusters. Sample sizes are the same as in J. **(L)** Same as in C but for GluA1-HaloTag. Examples of control (0 min) and cLTP (55 min). **(M)** Same as in D but for GluA1 nanoclusters. N=126-182 synapses from 9-13 neurons. **(N)** Same as in E but for GluA1 nanoclusters. Sample sizes are the same as in M. **(O)** Changes in normalized nanocluster number per PSD. Data are represented as mean ± s.e.m. N=10-18 neurons per group. **(P)** Changes in normalized nanocluster density. Data are represented as mean ± s.e.m. Sample sizes are the same as in O. **(Q)** Illustrative representation of the data shown in C-P. Top view shows the pictorial distribution of PSD95. Data are analysed using Kruskal-Wallis test. Post-hoc comparisons are done using Dunn’s test. \**p* < 0.05, \*\**p* < 0.01 and \*\*\**p* < 0.001.

The observed increase in the number of nanoclusters per PSD could be linked to an overall enlargement of the PSD. However, PSDs marked by PSD95 and GKAP did not expand up to 55 and 85 min after cLTP, respectively (Figure S2D). In contrast, we did observe an increase in the PSD area marked by SHANK3 5 min and 55 min after cLTP (Figure S2D). These data suggest that the size of the upper layers of the PSD does not significantly increase up to 55 min after cLTP, in accordance with previous studies.^17,18^

We next asked whether the increased density of PSD nanoclusters could be due to a decrease in nanocluster size. Indeed, the diameter of PSD95 and GluA1 nanoclusters was reduced 55 min after cLTP (Figure S2F). However, the diameter of GKAP and SHANK3 nanoclusters, which reside in deeper PSD layers, remained stable (Figure S2F). Thus, even though in the early phases of LTP the size of the PSD remains largely unaltered, synaptic activity drives rapid internal reorganization of scaffold nanoclustering within the PSD and causes repositioning of AMPA receptors into more tightly packed nanoclusters.

Lastly, we examined the alignment of PSD95 nanoclusters with the presynaptic scaffolding protein Munc13-1 (Figures S3A and S3B). We found that both the number of PSD95 and Munc13-1 nanoclusters increased after cLTP (Figures S3C and S3D), suggesting an increase in the number of release sites and transsynaptic nanocolumns. The fraction of PSDs in which Munc13-1 and PSD95 nanoclusters are aligned did not change after cLTP (Figures S3E and S3F), indicating that newly formed PSD95 nanoclusters are aligned with presynaptic release sites.

### Nanostructure of individual PSDs becomes more complex during LTP

Synapses are highly diverse in their structure, composition and strength. Thus, while we observe robust activity-induced changes in synaptic nano-organization on the population level, averaging over a large number of synapses could obscure the diversity in synaptic plasticity mechanisms. We therefore analysed the diversity in features of synapse nano-organization using hierarchical DBSCAN (HDBSCAN) and could distinguish four PSD types (Figures S4A and S4B). The majority of PSDs were classified as type 1 (62%) and had a simple morphology: average PSD size and one large nanocluster in the centre (Fig S4B, C). Type 2 PSDs are smaller and have two low-density nanoclusters (Figures S4B and S4C). Type 3 PSDs are the largest and have the highest complexity with on average three high-density nanoclusters (Figures S4B and S4C). Type 4 PSDs have an intermediate size and complexity, with two high-density nanoclusters (Figures S4B and S4C). We then estimated the relative occurrence of these PSD types before and after cLTP induction (Figure S4D). PSDs of type 1 reduced in number after cLTP induction from 62% to only 45% at 55 min. In contrast, PSD types 3 and 4 occurred more often after cLTP and increased from 5.8% to 13% and from 32% to 42%, respectively. This indicates that LTP increased the complexity in nano-organization of a subset of PSDs. Interestingly, type 2 PSDs were only observed directly after cLTP induction (1.4%), and possibly constitute newly formed, immature PSDs.

### Reorganization of AMPA receptor nanoclusters facilitates synaptic potentiation

We next asked whether the observed activity-induced reorganization of subsynaptic nanoclusters could be sufficient to potentiate synaptic transmission. We therefore developed a geometrical model of a canonical excitatory synapse that included a multistate activation scheme for AMPA receptors (Figure 2A). In a typical simulation, we measured the number of open AMPA receptors over time after the release of a single vesicle of glutamate, from which we also estimated the synaptic current (Figure 2B). To simulate nanoclusters with different densities, we placed AMPA receptors in clusters derived from normal distributions with varying standard deviations (*σ* = 30-70 nm; Figure 2C). Nanocluster diameter roughly corresponds to 2*σ*, such that cluster density is increased by decreasing the standard deviation of the normal distribution (Figure 2C). We then measured the maximum number of open receptors in synapses with an increasing number of AMPA receptors to simulate LTP. As expected, a higher total number of receptors led to an increase in receptor opening by glutamate release (Figure 2D). Interestingly however, the increase in the number of open receptors was higher for clusters with a low standard deviation (Figure 2D), indicating that clusters with higher density yield a stronger potentiation during the recruitment of AMPA receptors.

**Figure 2:**
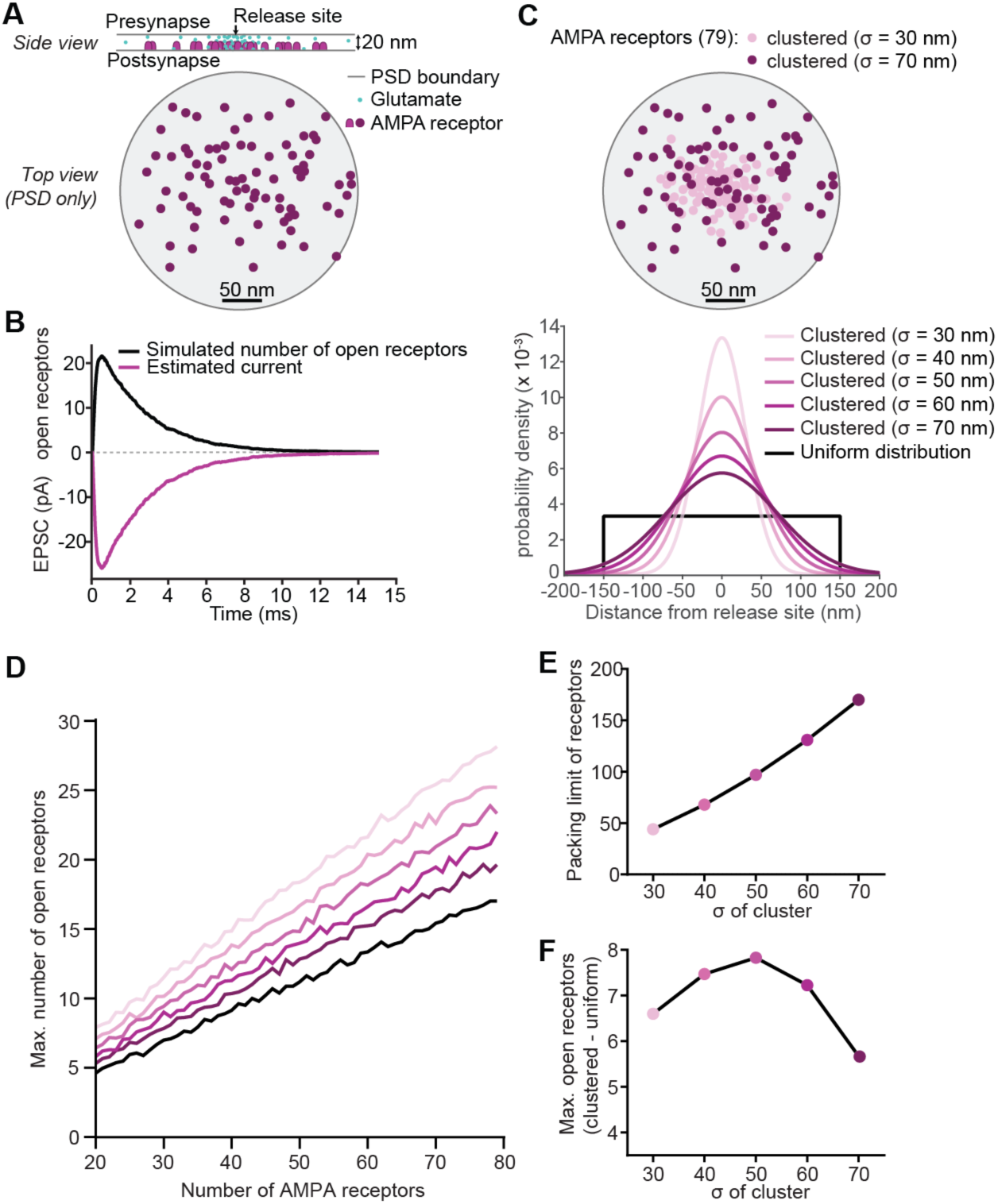
AMPA receptor nanoclustering can support synaptic potentiation. **(A)** Graphical representation of the MCell4 model of the synapse that was used for the Monte Carlo simulations. **(B)** Average number of open receptors (black line) and calculated EPSC (purple line) when 79 AMPA receptors were placed in a clustered distribution. **(C)** Top: Example AMPA receptor clusters based on normal distributions with σ = 30 nm and σ = 70 nm. Bottom: Normal distributions that were used to place clustered AMPA receptors in the model synapse. **(D)** The average maximum number of open AMPA receptors for all receptor distributions with increasing total receptor number. **(E)** The packing limit of AMPA receptors for the distributions used in D. **(G)** The difference in the maximum number of open AMPA receptors between the clustered and the uniform distribution, at the packing limit of the clustered distribution.

The AMPA receptors in these simulations are generally considered to be point-like, allowing for clusters with infinite density. However, AMPA receptor complexes have an approximated radius of ∼7.5 nm,^19,20^ limiting their maximum packing density in the synaptic membrane. We calculated the maximum packing density of AMPA receptors for different values of *σ* (Figure 2E). Then, we estimated for each packing limit how many extra AMPA receptors are opened in the clustered distribution compared to the uniform distribution to determine the maximum gain in synapse strength for each cluster size (Figure 2F). Remarkably, we found that the optimal standard deviation for clustering AMPA receptors close to a release site is ∼40-50 nm, corresponding to a nanocluster diameter of ∼80-100 nm. These results indicate that the maximum packing density of AMPA receptors limits the potentiating effect of nanocluster density on synapse strength and that the observed increase in nanoclusters (Figure 1) is necessary to sufficiently potentiate synaptic transmission.

### NMDA receptor activation is necessary for the LTP-induced increase in nanoclusters

Synaptic potentiation in the hippocampus is initiated by NMDA receptor activation and subsequent local calcium influx that triggers the activation of CaMKIIα. Therefore, we tested whether the nanoscale reorganization of the PSD that we observed after cLTP is dependent on NMDA receptors and CaMKIIα. We visualized endogenous PSD95 using HaloTag labelling and blocked either NMDA receptor currents using AP5 or prevented CaMKIIα activation using KN93 during cLTP induction (Figures 3A and 3B). The cultures were fixed directly or 55 min after cLTP induction and subsequently imaged using dSTORM. As before, the number and density of PSD95 nanoclusters did not change directly after cLTP (Figure 3C) but increased 55 min after cLTP (Figures 3B and 3D). This increase was prevented by AP5 treatment, while KN93 did not affect the increase in the number of PSD95 nanoclusters (Figure 3D). Surprisingly however, the cLTP-induced increase in nanocluster density was not blocked by AP5 and only partially by KN93 (Figure 3D). In summary, NMDA receptor activation is necessary for the increase in PSD95 nanoclusters during LTP but is not required for the increase in nanocluster density. Conversely, CaMKIIα activation is not necessary for the cLTP-induced increase in nanoclusters but seems at least partially required to create more tightly packed PSD95 nanoclusters.

**Figure 3:**
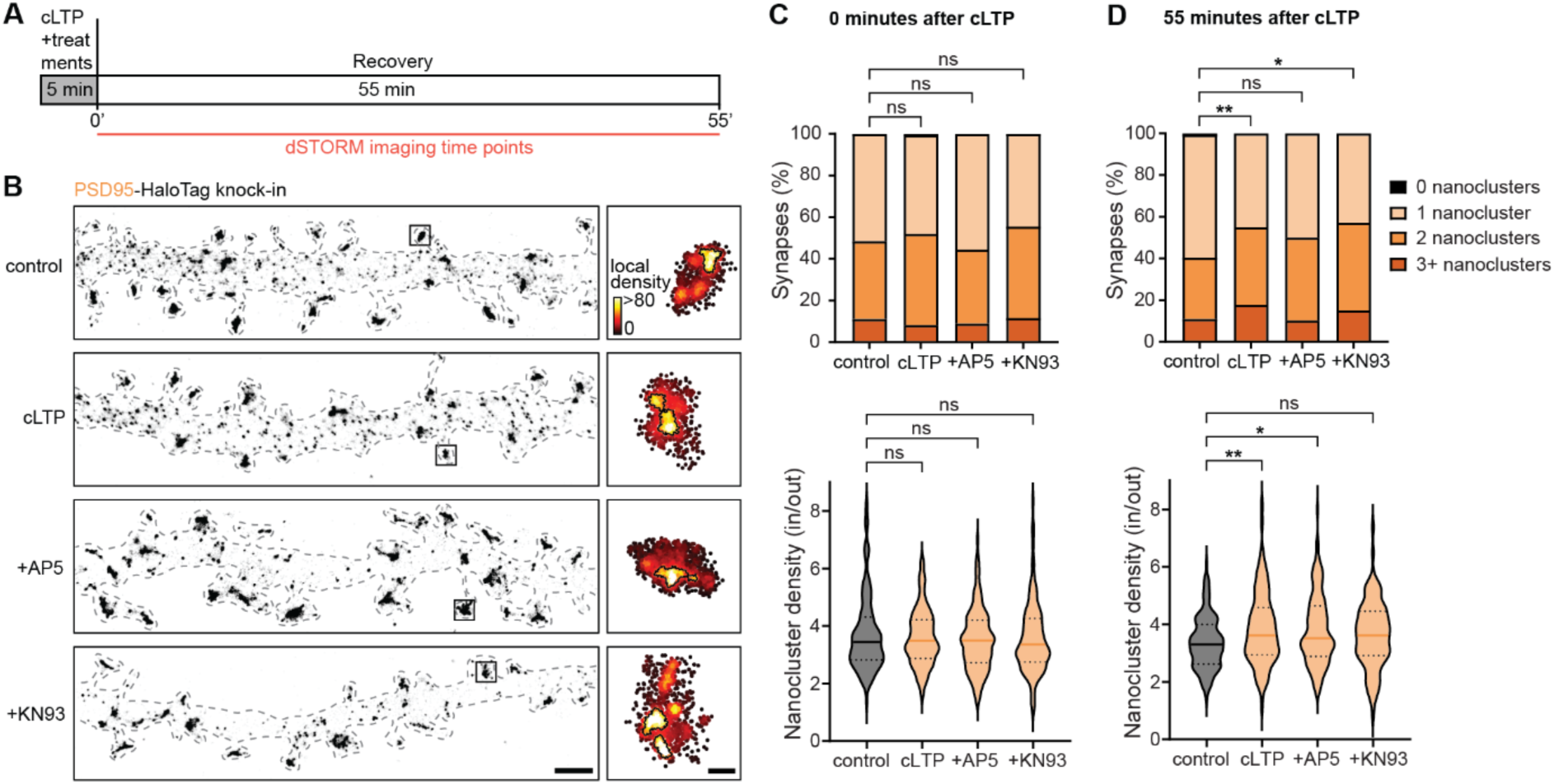
cLTP-induced increase in PSD95 nanoclusters is dependent on NMDA receptor activation. **(A)** Timeline of cLTP experiment. AP5 and KN93 were included in the cLTP medium only. **(B)** Left: Representative dendritic stretches of reconstructed dSTORM images of PSD95-HaloTag labelled with JF646, 55 min after cLTP induction for the conditions control, cLTP, +AP5 and +KN93. Scale bar: 1 µm. Right: PSDs indicated with black squares are shown after nanocluster analysis. Scale bar: 200 nm. **(C)** Number of PSD95 nanoclusters per PSD and nanocluster density at 0 min. Top: Stacked histogram of nanocluster number per PSD. Bottom: Violin plot of the relative density of PSD95 nanoclusters. Sample size: N=130-150 synapses from 13-15 neurons. **(D)** Same as in C but for the time point 55 min after cLTP induction.. Sample sizes are the same as in C. Data are analysed using Kruskal-Wallis test. Post-hoc comparisons are done using Dunn’s test. \**p* < 0.05, \*\**p* < 0.01 and \*\*\**p* < 0.001.

### Actin remodelling is required for PSD reorganization during LTP

It remains unclear what molecular mechanisms drive the internal reorganization of scaffolding proteins. Actin regulators are dynamically recruited during LTP to remodel the spine actin cytoskeleton,^17^ which could in addition restructure the PSD. Indeed, depolymerization of the actin cytoskeleton leads to fewer and smaller PSD95 nanoclusters,^1^ whereas treadmilling of F-actin can reposition AMPA receptors inside the PSD.^21^ We thus hypothesized that actin treadmilling and remodelling of the actin cytoskeleton drive the cLTP-induced internal reorganization of the PSD. Jasplakinolide treatment during cLTP (stabilizing F-actin polymers) prevented cLTP-induced formation of high-density nanoclusters of PSD95 (Figure S5). Unexpectedly, brief latrunculin B treatment (depolymerising F-actin polymers) with subsequent washout increased the number and density of nanoclusters and seemed to occlude the cLTP-induced reorganization of PSD95 nanoclusters (Figure S5). Thus, interfering with actin dynamics during LTP induction prohibits activity-induced reorganization of the PSD.

### The SHANK protein family of scaffolding molecules is essential for plasticity-induced reorganization of the PSD

We next asked how F-actin dynamics can drive the activity-induced nanoscale reorganization of the PSD. The SHANK family of proteins can bind multiple other PSD scaffolding proteins and interact with the actin cytoskeleton, which makes them ideally positioned to mediate internal reorganization of the PSD. ^11–14^ In addition, we observed that the number of SHANK3 nanoclusters increased immediately after cLTP induction (Figure 1O), preceding the reorganization of other PSD scaffolds. To test whether SHANK proteins are involved in cLTP-induced PSD reorganization, we used a previously validated miRNA-based knockdown construct that simultaneously depletes the three SHANK1-3 paralogs expressed at excitatory synapses (Figures S5A and S5B).^14^ We found that the cLTP-induced increase in PSD95 nanocluster number and density was completely absent in SHANK1-3 knock-down neurons (Figures S5C-E), and fully rescued by re-expression of SHANK3 (Figure 4). On the other hand, while SHANK2 re-expression partially restored the increased in nanocluster number, this was not sufficient to fully rescue the cLTP-induced reorganization of the PSD (Figure 4). Thus, activity-induced internal reorganization of the PSD is dependent on SHANK scaffolding proteins, particularly SHANK3.

**Figure 4:**
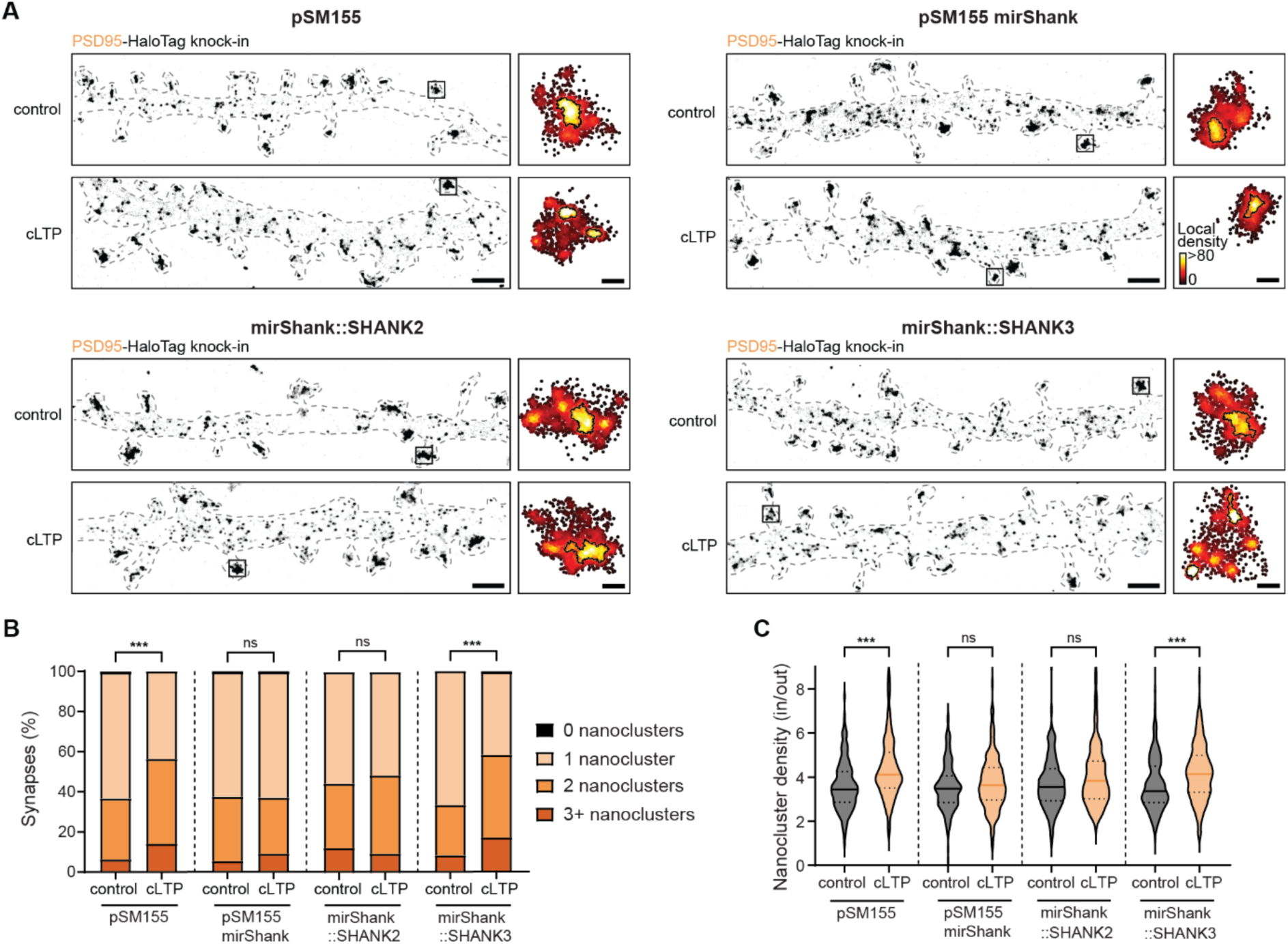
SHANK3, but not SHANK2, is required for cLTP-induced internal reorganization of the PSD. **(A)** Shown are representative dendritic stretches of dissociated hippocampal neurons with highlighted PSDs 55 min after cLTP induction. Scale bar left: 1 µm. Scale bar right: 200 nm. **(B)** PSD95 nanocluster number 55 min after cLTP induction.. Sample size: N=136-221 synapses from 8-13 neurons **(C)** Violin plot of the relative density of PSD95 nanoclusters. Sample sizes are the same as in B. Data are analysed using Kruskal-Wallis test. Post-hoc comparisons are done using Dunn’s test. \**p* < 0.05, \*\**p* < 0.01 and \*\*\**p* < 0.001.

### SHANK3-actin interactions mediate LTP-induced PSD reorganization

To determine which protein-protein interactions drive SHANK3-mediated PSD reorganization, we developed mutant versions of SHANK3 that decouple the upper PSD layers from the actin cytoskeleton but leave synapse size and number intact (Figure 5A). SHANK3 is coupled to PSD95 and the upper layers of the PSD via GKAP.^10,22^ To test the necessity of GKAP-SHANK3 interactions, we reduced SHANK3-GKAP binding by deleting a sequence close to the PDZ domain of SHANK3 (Figure 5A; ΔT543-Y564), which is known to reduce the affinity of SHANK3 for GKAP ∼5 times.^23^ SHANK3ΔT543-Y564 was still highly enriched in PSDs and this deletion did not affect synapse number (Figures 5B, S6 and S6B). The ability of SHANK3, but not SHANK2, to rescue PSD reorganization suggests that perhaps interaction domains specific to SHANK3 mediate changes in synaptic nano-organization. Notably, recently it was found that the N-terminal SPN/ANK domains that are unique for SHANK3 interact with F-actin in a regulated manner.^12^ To test whether this interaction is required for cLTP-induced PSD reorganization, we developed a minimal mutant with two missense mutations that are known to diminish SHANK3 binding to F-actin (Figure 5A; Q37A/R38A).^12^ Importantly, SHANK3-Q37A/R38A localized to PSDs and the mutation did not affect synapse number (Figures 5B, S6C and S6D).

**Figure 5:**
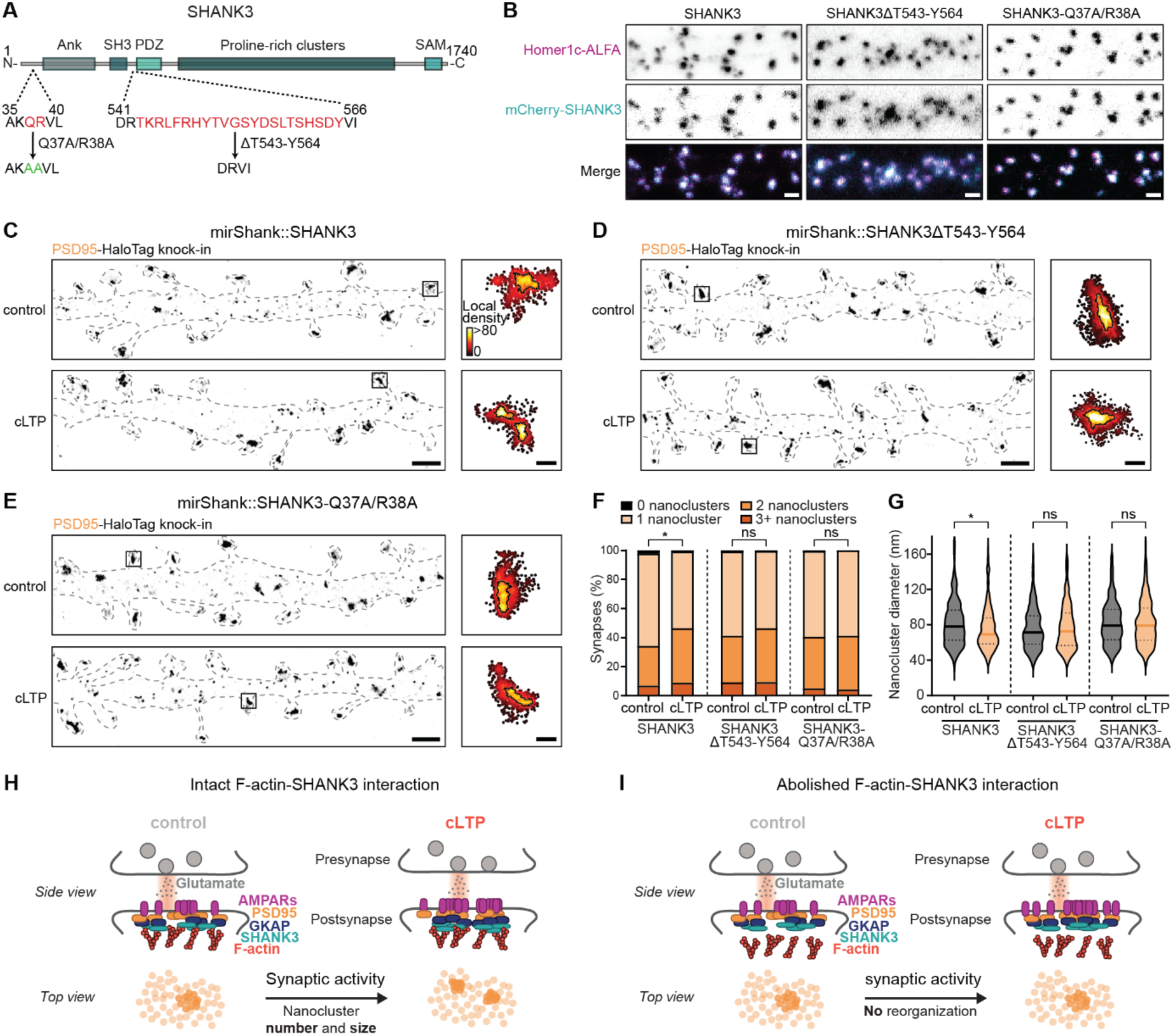
Direct F-actin-SHANK3 interactions are necessary for internal PSD reorganization during LTP. **(A)** Schematic overview of the SHANK3 rescue construct and the ΔT543-Y564 deletion and the Q37A/R38A mutation. **(B)** Example confocal images of dendritic stretches of dissociated hippocampal neurons. Scale bar: 2 µm. (**C-E)** Left: Representative dendritic stretches 55 min after cLTP induction for the conditions SHANK3 (C), SHANK3ΔT543-Y564 (D) and SHANK3-Q37A/R38A (E). Scale bar: 1 µm. Right: PSDs indicated with black squares are shown after nanocluster analysis. Scale bar: 200 nm. **(F)** PSD95 nanocluster number per PSD 55 min after cLTP induction. Sample size: N=209-304 synapses from 11-16 neurons **(G)** Violin plot of PSD95 nanocluster diameter 55 min after cLTP induction. Sample sizes are the same as in F. **(H)** Illustrative representation of an excitatory synapse before and after cLTP induction, when F-actin-SHANK3 interaction is intact. Top view shows pictorial distribution of PSD95. **(I)** Same as in H but in the case of abolished F-actin-SHANK3 interaction.

Confirming our earlier results, re-expression of wildtype SHANK3 rescued the cLTP-induced increase in PSD95 nanoclusters (Figures 5C and 5F). However, re-expression of SHANK3ΔT543-Y564, reducing the SHANK3-GKAP interaction, did not completely rescue the increase in nanoclusters (Figures 5D and 5F), whereas re-expression of SHANK3-Q37A/R38A, interfering with the SHANK3-actin interaction, completely prevented cLTP-induced PSD95 reorganization (Figures 5E and 5F). SHANK3 wildtype re-expression also led to a reduction in nanocluster diameter after cLTP (Figure 5G), which did not occur when SHANK3ΔT543-Y564 and SHANK3-Q37A/R38A were expressed. Nanocluster density did not change significantly (Figure S6E). In conclusion, diminishing F-actin-SHANK3 interactions completely abolished the cLTP-induced nanoscale reorganization of the PSD, whereas reducing SHANK3-GKAP binding attenuated the increase in nanocluster number. These results indicate that activity-induced actin dynamics can drive the intricate nanoscale reorganization of the PSD through interactions with SHANK scaffolding proteins.

## Discussion

In this study, we resolved the activity-induced nanoscale reorganization of endogenous GluA1, PSD95, GKAP and SHANK3. Intriguingly, this activity-induced reorganization originated in deep PSD layers and relied on the interaction between SHANK3 and the actin cytoskeleton. Furthermore, our computational simulations demonstrate that the observed increase in nanocluster density and number is sufficient to facilitate synaptic potentiation. Together, our results reveal a novel molecular framework for how nano-structural remodelling of postsynaptic protein assemblies is regulated to support synaptic plasticity.

A key finding of this study is that an almost immediate internal reorganization of PSD scaffold nanoclusters precedes the increase in the number of AMPA receptor nanoclusters. This delayed formation of AMPA receptor nanoclusters suggests that AMPA receptors newly inserted into the synaptic membrane preferentially populate pre-formed scaffold nanoclusters. The computational simulations show that increasing AMPA receptor nanocluster density potentiates postsynaptic responses, consistent with previous studies.^1,3^ Importantly however, our simulations also indicate that when existing nanoclusters reach their packing limit, formation of new nanoclusters is required to maximize the number of clustered receptors. Nevertheless, formation of new nanoclusters will only potentiate synaptic strength if the postsynaptic nanocluster resides close to the presynaptic release site. Indeed, we found that PSD95 nanoclusters align with release sites after LTP, suggesting that the majority of newly formed nanoclusters reside near a presynaptic release site. Our findings thus reveal that the internal nanoscale reorganization of PSD scaffold proteins is a precisely orchestrated process that is essential to facilitate subsequent receptor positioning and potentiate synaptic strength.

Our study defines a critical role for SHANK3 in the activity-induced nanoscale reorganization of the PSD. We found that SHANK3 nanoclusters were the first to increase after LTP induction and that SHANK3 is necessary for the activity-induced increase in PSD95 nanocluster number and density. Consistent with a role for SHANK3 in reorganizing PSD organization during LTP, complete loss of SHANK3 was shown to alter PSD95 nanostructure *in situ* and to lead to deficits in LTP.^24–26^ Our knockdown-rescue studies further established that interactions between SHANK3 and the actin cytoskeleton, and between SHANK3 and GKAP scaffolds, are required for activity-induced PSD reorganization. These findings support a model where SHANK proteins integrate forces generated by activity-induced actin cytoskeleton polymerization and branching and propagates these forces through scaffold-scaffold interactions to the postsynaptic membrane. Liquid-liquid phase separation (LLPS) mechanisms could further support activity-dependent remodelling of SHANK3 and other PSD components.^27–31^

Spine actin dynamics drive ongoing and activity-dependent changes in spine and PSD morphology^14,17,21,32^ and depolymerization of the actin cytoskeleton acutely disrupts the maintenance of scaffolding nanoclusters as well as the subsynaptic distribution of AMPA receptors.^1,21,33^ Here we significantly extend these findings, demonstrating that activity-induced changes in actin dynamics control reorganization of the PSD directly through SHANK proteins. Apart from direct interactions with the actin cytoskeleton, SHANK proteins can interact with several actin nucleation factors such as Cortactin, Abp1, the WAVE and the Arp2/3 complex.^10,11,13,34^ Furthermore, previous super-resolution studies demonstrated that foci of high actin polymerization localize preferentially near the PSD^35^ and that the Arp2/3 and WAVE complex are closely associated with the PSD.^36^ In addition, SHANK proteins can recruit a plethora of signalling proteins such as CaMKII,^27,28,37^ HRas and Rap1,^12,38^ as well as modulators of the small Rho/Rac/Cdc42 GTPases that are involved in regulating spine actin polymerization and branching.^39–41^ SHANK nanoclusters could thus serve as core scaffolding platforms orchestrating localized signalling events that act to drive focal actin polymerization and nucleation to exert directed forces on the PSD. Future studies will need to determine the spatiotemporal dynamics governing the association of these components with SHANK nanoclusters.

Altogether, our findings provide a new structural framework for how activity-induced actin dynamics control postsynaptic reorganization. Even broader, we believe our findings illuminate a new specific model of how the cytoskeleton dynamically interacts with and modifies multiprotein complexes to drive cellular signalling. Importantly, since SHANK mutations are strongly associated with Autism Spectrum Disorder (ASD),^47^ and particularly ASD mutations in SHANK3 impair association with actin,^42^ investigating the effects of ASD mutations on synapse nano-organization could help elucidate how these mutations disrupt synaptic function.

## Methods

### Animals

All animal experiments were performed in compliance with the guidelines for the welfare of experimental animals issued by the Federal Government of The Netherlands and were approved by the Dutch Central Committee Animal experiments, Project AVD10800202216383. Timed pregnant Wistar rats were obtained from Janvier labs for dissociated neuronal cultures.

### Molecular cloning

All vectors that were used in this study are listed in Table S1. All oligonucleotides used to make these vectors are listed in Table S2. pORANGE *Dlg4*-HaloTag was based on pORANGE *Dlg4*-GFP (Addgene #131477),^16^ where GFP was replaced with HaloTag using HindIII and XhoI, and PCR of oligos Oligo01 and Oligo02. pORANGE *Gria1*-HaloTag was based on pORANGE *Gria1*-GFP (Addgene #131489),^16^ were GFP was replaced with HaloTag using BmtI and MluI, and PCR of oligos Oligo03 and Oligo04. pOC2 *Dlgap1*-HaloTag was made in two steps. First, oligos Oligo05 and Oligo06 were inserted in pOC2 cloning template vector (Addgene #183421)^15^ using BbsI and primer ligation. Then, the donor DNA with HaloTag was made using PCR of oligos Oligo07 and Oligo08 and ligated in HindIII and MluI sites of the intermediate construct. pOC4 FLAG-Cas9 (Addgene #183423) was described before.^15^ pX333 *Shank3*-HaloTag was made by two primer ligations. Firstly, oligos Oligo09 and Oligo10 were ligated in the Eco31I site of pX333 (Addgene #64073, a generous gift from Andrea Ventura),^43^ to express a gRNA with donor recognition sequence 3 (DRS-3) from Gao et al. (2019).^44^ Secondly, oligos oWD0219 and oWD0220 were ligated in the BbsI site of pX333, to express a gRNA with the target sequence of *Shank3*. For the same *Shank3*-HaloTag knock-in, we first made the donor DNA with HaloTag using PCR of oligos oWD0217 and oWD0218, and ligated it in the EcoRV site of pJet1.2 (Thermo Fisher Scientific). Secondly, we replaced HaloTag with HaloTag-STOP which was isolated from pORANGE *Gria1*-HaloTag using NheI and AfeI.

pCaMK Homer1c-mCherry was described before.^45^ pCaMK Homer1c-mRuby3 was created based on pCaMK Homer1c-mCherry using HiFi assembly. pCaMK Homer1c-ALFA was made by digestion of pCaMK Homer1c-mCherry with BshTI and NotI, and primer ligation of oligos Oligo13 and Oligo14.

pSM155 mCherry, pSM155 mirShank::mCherry, pSM155 mirShank::mCherry-SHANK2 and pSM155 mirShank::mCherry-SHANK3 were previously described.^14,46^ pSM155 mirShank::mCherry-SHANK3ΔT543-Y564 was made by site-directed mutagenesis using in vivo assembly (IVA),^47^ with oligos Oligo15 and Oligo16. pSM155 mirShank::mCherry-SHANK3-Q37A/R38A was made by site-directed mutagenesis using IVA with oligos Oligo17 and Oligo18. All plasmids created in this study will be made available on Addgene.

### Dissociated neuronal cultures and transfection

Hippocampi of embryonic day 18 rat brains from both genders were dissociated using trypsin and plated on Ø18-mm coverslips (type 1.5) coated with poly-L-lysine (37.5 µg/ml, Sigma Aldrich) and laminin (1.25 µg/ml, Roche Diagnostics) at a density of 100,000 neurons per well. Neurons were cultured in Neurobasal (NB) with 0.5 mM L-glutamine, 1% penicillin + streptomycin (pen/strep) and 2% B27 (NB-complete, all Invitrogen) until day in vitro (DIV) 7, after which the medium was refreshed every 7 days by replacing 400 µl medium with 500 µl BrainPhys (BP) supplemented with 2% NeuroCult SM1 (Stem Cell Technologies) and 1% pen-strep. Dissociated cultures were incubated at 37°C in 5% CO_2_.

Dissociated hippocampal neurons were transfected at DIV 3-4 using Lipofectamine 2000 (Invitrogen). Per coverslip, 3.3 µl of Lipofectamine was mixed with DNA vectors in 200 µl NB and incubated for 30 min at RT. To start transfection, 500 µl of conditioned medium was replaced by the 200 µl DNA/Lipofectamine mix. To increase transfection efficiency, an extra 200 µl of conditioned medium was removed. Neurons were incubated for 90-120 min at 37°C in 5% CO_2_. Then, coverslips were placed in new wells containing 500 µl conditioned medium and 500 µl NB-complete and maintained at 37°C in 5% CO_2_ until the day of the experiment.

### Chemical LTP (cLTP)

For cLTP experiments, extracellular media E4 (only Figures 1C-E and 3) and E5 were used. E4 medium consisted of (in mM) 120 NaCl, 3 KCl, 10 HEPES, 2.7 CaCl_2_, 2 MgCl_2_, and 10 D-glucose (pH set to 7.3-7.4). E5 medium consisted of (in mM) 140 NaCl, 3 KCl, 10 HEPES, 2.7 CaCl_2_, 2 MgCl2 and 10 D-glucose, and was adjusted to an osmolarity of 315-320 mOsm using sucrose (pH set to 7.3-7.4). cLTP medium was based on these extracellular media but did not contain MgCl_2_ and was supplemented with 25 µM bicuculline and 260 µM glycine. To induce cLTP, coverslips with dissociated rat hippocampal neurons (DIV 21-23) were transferred to wells containing 500 µl E4 / E5 (control) or cLTP medium and were incubated for 5 min at 37°C. Depending on the experimental condition, neurons were then either fixed or placed in 500 µl E4 / E5 and incubated at 37°C to recover until fixation. For experiments that are depicted in Figures 1I-N and 5C-G, dissociated hippocampal neurons were pre-incubated for 20 min in E5 at 37°C and 5% CO_2_ before cLTP induction.

### Immunohistochemistry

Dissociated neuronal cultures were fixed in PBS with 4% paraformaldehyde (PFA) and 4% sucrose for 5 min at RT. If cultures were subsequentially used for dSTORM, neurons were fixed in PEM80 buffer consisting of 80 mM PIPES, 2 mM MgCl_2_, and 1 mM EGTA (pH 6.8), with 4% EM-grade PFA (Electron Microscopy Sciences) for 5 min at 37°C. Next, neurons were washed three times in PBS with 0.1 M glycine (PBS/gly) and blocked in PBS/gly supplemented with 10% (v/v) normal goat serum (NGS, Abcam) and 0.1% (v/v) Triton X-100 for 30 - 60 min at 37°C. If the experiment required a cell surface-enriched staining of the epitope, Triton X-100 was excluded from blocking and antibody solutions. After blocking, primary antibodies were diluted in PBS/gly with 5% (v/v) NGS and 0.1% (v/v) Triton X-100 and incubated with the neurons for 2 hours at RT or overnight at 4°C. After being washed three times in PBS/gly, neurons were incubated with secondary antibodies diluted (1:400 -800) in PBS/gly with 5% (v/v) NGS and 0.1% (v/v) Triton X-100 for 1 hour at RT. Finally, neurons were washed three times in PBS/gly, dipped in milliQ water (MQ) and mounted in polyvinyl alcohol mounting medium with DABCO (Supelco), or Fluoromount-G (Invitrogen). All antibodies and reagents used in this study can be found in Table S3.

### Live-cell HaloTag labelling

Constructs fused to HaloTag were live-labelled with HaloLigand conjugated to Janelia Fluor dyes JFX554 (a generous gift from Dr. Lavis, HHMI Janelia Research Campus) or JF646 (Promega), maximally 20 min before the experiment. Neurons were incubated with the HaloLigand (0.2 µM in conditioned medium) for 10 min at 37°C in 5% CO_2_ and then returned to the conditioned medium to recover.

### Confocal microscopy

Confocal images were acquired with a Zeiss LSM700 or LSM900 microscope equipped with 405-nm, 488-nm, 561-nm, and 642-nm excitation lasers. All confocal images of dissociated rat hippocampal neurons are presented as maximum-intensity projections of z-stacks consisting of 3 -5 imaging planes with 0.5-µm intervals, pixel size of 0.1 µm, and averaged 2x. A Plan-Apochromat 63x oil (N.A. 1.40) objective was used to acquire the images. Zoom 0.6x was used for confocal images presented in Figure S1.

### Single-molecule localization microscopy

SMLM experiments dSTORM were carried out on the Nanoimager S from Oxford Nanoimaging Ltd. (ONI). The ONI Nanoimager is equipped with a 100x oil-immersion objective (N.A. 1.40), four laser lines (405 nm, 473 nm, 561 nm, and 640 nm), a sCMOS camera (ORCA Flash 4, Hamamatsu) and a closed-loop piezo stage. An integrated filter set splits the blue, green, and red emission spectra to the left side of the camera, and far-red emission to the right side. The temperature of the microscope box, stage, and objective was kept constant at 30°C to minimize drift during acquisitions. The drift in z was corrected during imaging using the near-infrared auto-focus laser (720 nm). Drift in x and y was corrected post-imaging in the NimOS software or using the Detection of Molecules (DoM, v.1.2.5) plugin in ImageJ (https://github.com/UU-cellbiology/DoM_Utrecht). During imaging, coverslips with dissociated rat hippocampal neurons were vacuum sealed in dSTORM buffer consisting of PBS, 5% w/v D-glucose, 40 µg/ml catalase (Sigma-Aldrich), 700 µg/ml glucose oxidase (Sigma-Aldrich) and 5-35 mM cysteamine (MEA). dSTORM acquisitions were acquired at 50 Hz using an exposure time of 20 ms.

### Clustering PSD nano-organizations using HDBSCAN

All PSD95 dSTORM data that was acquired and analysed using the nanocluster analysis was pooled to unravel PSD types based on their nano-organization. PSDs were excluded if they lacked a nanocluster. In total 40,279 PSDs were included and the variables PSD diameter (µm), nanoclusters per area PSD (µm^-2^), nanocluster density (in/out), nanocluster diameter (µm), and distance between nanocluster and PSD centre (µm) were used to identify PSD types.

First, in Python 3.11, we used Box-Cox transformation and the function *boxcox()* from the *sklearn.preprocessing* module to reduce the heteroscedasticity of the variables. Second, we scaled all data to centre each variable at zero using *StandardScaler()* from the *scipy.stats* module. Then, we clustered all PSDs using HDBSCAN using *HDBSCAN()* from the *hdbscan* module, with min_samples=8, min_cluster_size=7, cluster_selection_epsilon=0.4, cluster_selection_method=’leaf’, alpha=1.0, leaf-size=5. PSDs were clustered in four different types. Type 1 contained 21,046 PSDs, type 2 only 44, type 3 had 2,409 PSDs, type 4 had 9,981 PSDs, and 6800 PSDs were not considered to be part of a cluster by the HDBSCAN algorithm. We utilized Uniform Manifold Approximation and Projection (UMAP) to visualize the clusters of PSD types in two dimensions (Figure S4). In Python 3.11, *UMAP()* from the *umap* module was used to reduce the number of dimensions, with n_neighbors=15, min_dist=0.0, n_components=2, metric=’eaclidean’, random_state=100. The Python code used for this analysis (DOI: 10.5281/zenodo.11098185) and the PSD95 dataset can be found in the GitLab repository at https://gitlab.com/maclab/hdbscan-of-psd-nanostructure.

### Monte Carlo simulations of synaptic transmission

A geometrical model of excitatory synapses was constructed in CellBlender using <u>MCell4</u>. The model consisted of a pre- and postsynaptic circular plane with a diameter of 1 µm separated by 20 nm. AMPA receptors were placed inside a designated PSD area that was 300 nm in diameter. AMPA receptors were 15 nm in diameter,^19,20^ had a diffusion coefficient of 1 nm^2^ / ms and had six closed states and one open state.^48^ The AMPA receptor state followed the scheme below:

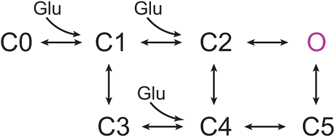

and reaction rates are described in Table S4. Each iteration of the Monte Carlo simulations lasted for 15 ms and had a time step of 1 µs. The simulation started with the release of 8,000 glutamate molecules from a point source in the middle of the presynaptic plane. Glutamate molecules diffused with a diffusion coefficient of 0.4 µm^2^ / ms and could escape through the edge of the synaptic cleft (reaction rate = 2.5 · 10^3^).^49^ Upon interaction with glutamate, AMPA receptors could switch from one state to another depending on the reaction rate (Table S4). During a simulation, the number of open receptors was scored for each time step. To determine the EPSC amplitude and shape, the number of open receptors was multiplied by a current of -1.2 pA, assuming a membrane potential of -65 mV and a single channel conductance of 18.8 pS.^50^

A multivariate normal distribution function (MVN) was used to determine the placement of AMPA receptors in the synapse:

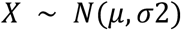

where *μ* is the centre and *σ* is the standard deviation of a normal distribution. AMPA receptor coordinates were then drawn from this normal distribution. Each iteration drew new coordinates from the same distribution. These distributions were used to model the density of nanoclusters, in which a value of *σ* corresponds roughly to the radius of a nanocluster. A total of 200 iterations where performed for each value of *σ*. The packing limit of each cluster was calculated using a receptor width of 15 nm.^19,20^

All simulations were performed on a desktop computer running 64-bit Ubuntu 20.04.4 LTS (GNOME v3.36.8), with an Intel HD Graphics 4600 (HSW GT2) graphics card, an Intel Core i5-4670 CPU @ 3.40GHz × 4 processor, and 15.5 Gb RAM. The MCell4 model of the synapse including all scripts necessary to reproduce the simulations (DOI: 10.5281/zenodo.11098504) can be found in the GitLab repository at https://gitlab.com/maclab/mcell4-synapse-model.

## Quantification and analysis

### Synapse number and fluorescence intensity

For experiments depicted in Figures 5B, S2A, B and S6A-D, the ComDet (v0.5.5) plugin in ImageJ (https://github.com/UU-cellbiology/ComDet) was used to identify synapses based on Homer1c-ALFA fluorescence signal. The intensity threshold to identify puncta in the maximum-intensity projection of confocal images of Homer1c-ALFA was 12 (Figures 5B and S6A-D) or 17 (Figures S2A and S2B). Synaptic intensity was measured as the total fluorescence intensity inside the detected synapse puncta (*S_in_*) subtracted by the average fluorescence intensity in the perimeter of the bounding rectangle (*I_bg_*) multiplied by the number of pixels inside the puncta (*N*). Synaptic enrichment was calculated as the synaptic intensity divided by the fluorescence intensity in the dendritic shaft.

To measure surface levels of SEP-GluA1 (Figures S2A, and S2B), an ROI was drawn around a representative dendrite of the maximum-intensity projection image of each cell in ImageJ based on the fluorescence signal of SEP-GluA1. Then, the fluorescence intensity of Alexa Fluor 647 in that ROI was measured to determine the surface levels of SEP-GluA1.

### Nanocluster analysis of SMLM data

The NimOS software from ONI (v18.4 and v19.7) was used to detect single-molecule localization events. Subsequently, MATLAB (v2021a) was used to filter the localizations and resolve nanoclusters inside PSDs. Localizations were included if they met the following criteria: standard deviation of localization error <15 nm (exceptions: <20 nm for SHANK3 and GluA1), localization persisted for <10 frames, and the number of cumulative photons of localizations was >300 and <30,000. ROIs encircling PSDs were based on the fluorescence signal of Homer1c or PSD95, automatically detected by thresholding the signal intensity and outlined using the full-width half maximum (full-width quarter maximum for GluA1) around the fluorescence peak of each PSD. PSDs were accepted for nanocluster analysis if they contained more than 800 localizations and if their size was > 0.02 µm^2^ and <0.3 µm^2^ (<0.4 µm^2^ for GluA1).

Nanocluster analysis was based on.^1^ A localization was considered to be part of a nanocluster if its local density (= number of other localizations within a radius of 5x the mean nearest neighbour distance) was >40. Nanoclusters were then identified using the functions *linkage()* and *cluster()*. Nanoclusters were subdivided if they contained multiple density peaks separated by local minima of <30% of the highest density peak. Boundaries of nanoclusters were constructed using Voronoi diagrams encompassing the outer localizations. Nanoclusters were only included if they comprised >5% of the localizations (in that PSD), and if they were >30 nm in diameter. PSD size and synaptic area (for GluA1) were calculated based on the cluster of localizations inside the ROI, which was determined using DBSCAN with epsilon = 5x mean nearest neighbour distance and minpts = 4.^51^

After the nanocluster analysis, each dataset was balanced (except data in Figure S5) by selecting a random subset of analysed synapses from each acquisition. The mean measures of this subset did not differ more than 5% from the means of the whole acquisition. For each dataset, the number of selected synapses per acquisition was equal to the lowest number of acquired synapses in that particular dataset. MATLAB scripts used for nanocluster analysis (DOI: 10.5281/zenodo.11098540) can be found in the GitLab repository at https://gitlab.com/maclab/nanocluster-analysis-of-smlm.

### Statistics

At least four independent dissociated rat hippocampal cultures were used for each experiment. When two groups were compared, an unpaired *t* test was used. If the groups were not normally distributed, the non-parametric Mann-Whitney *U* test was used. A one-way ANOVA was used to compare multiple groups if they were normally distributed and had equal variances. A Holm-Šídák test was used for *post hoc* correction and comparisons. If the groups were not normally distributed or did not have equal variances, a Kruskal-Wallis test was used. In that case, Dunn’s test was used for *post hoc* correction and comparisons. All statistics were performed in GraphPad Prism 10.

## Acknowledgements

This work was supported by the Nederlandse Organisatie voor Wetenschappelijk Onderzoek (NWO) Grant OCENW.KLEIN.163 (to H.D.M.). We kindly thank Jelmer Willems and Manon Westra for their contributions to the MATLAB code used for nanocluster analysis. We want to thank the whole MacGillavry lab for thorough discussions and suggestions.

## Author contributions

Conceptualization, W.J.D., F.B. and H.D.M.; methodology, W.J.D., N.R.R., F.B. and H.D.M.; investigation, W.J.D., A.K.S., N.R.R., L.C.P.S, A.A.M. and O.K.K.; analysis, W.J.D., A.K.S., N.R.R., L.C.P.S, A.A.M. and O.K.K; writing original draft, W.J.D. and H.D.M.; editing manuscript, W.J.D., F.B. and H.D.M.; visualization, W.J.D.; supervision, F.B. and H.D.M.; funding acquisition, H.D.M.

## Competing interests

The authors declare no competing interests.

## Materials and correspondence

Requests for further information and resources should be directed to the lead contact Harold MacGillavry.

## Data availability

All source and raw data will be deposited on Yoda upon publication under a CC BY 4.0 license for open data sharing.

## Computer code

The analyses and scripts used for nanocluster analysis (DOI: 10.5281/zenodo.11098540), MCell4 simulations of synaptic transmission (DOI: 10.5281/zenodo.11098504) and HDBSCAN clustering of PSD nanostructure types (DOI: 10.5281/zenodo.11098185) can be found at https://gitlab.com/maclab.

---

**Figure S1:**
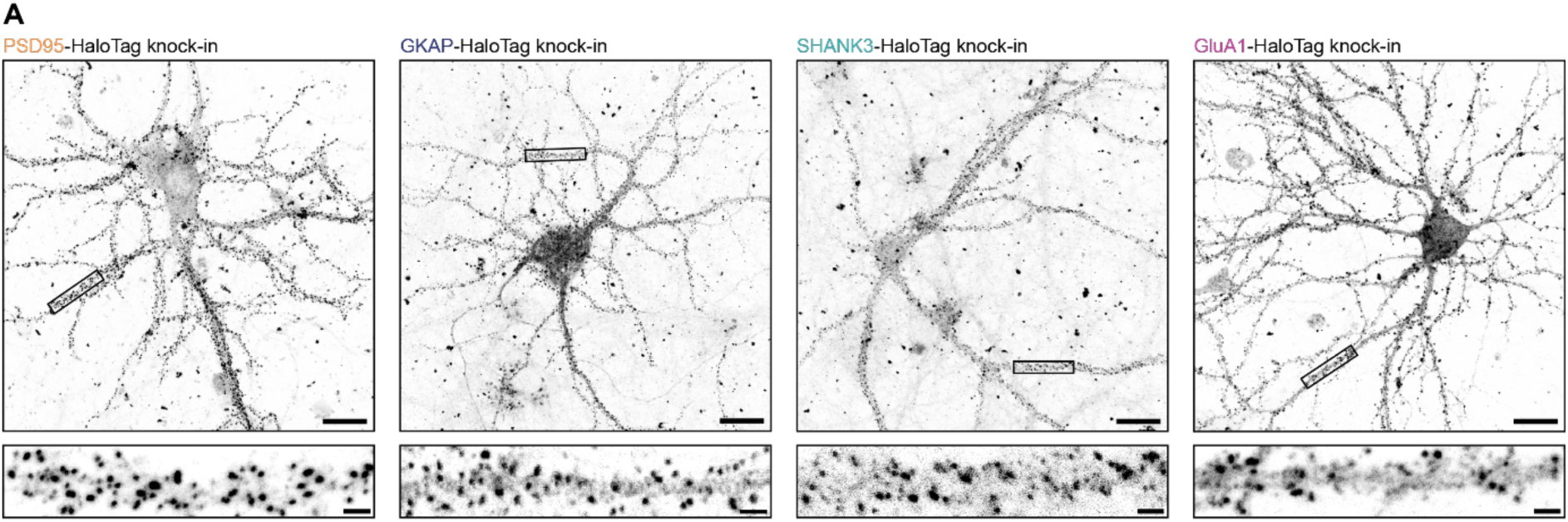
ORANGE knock-ins for tagging endogenous PSD95, GKAP, SHANK3 and GluA1 in hippocampal neurons. **(A)** CRISPR/Cas9-mediated knock-ins were created to tag PSD95, GKAP, SHANK3 and GluA1 endogenously (Table S1). Dissociated rat hippocampal neurons were transfected at DIV 4 and fixed at DIV 21. HaloTag was labelled with JFX554 and visualized using confocal microscopy. Shown images are maximum-intensity projections of the acquired z stacks. Scale bar top images: 20 µm. Scale bar bottom images: 2 µm.

**Figure S2:**
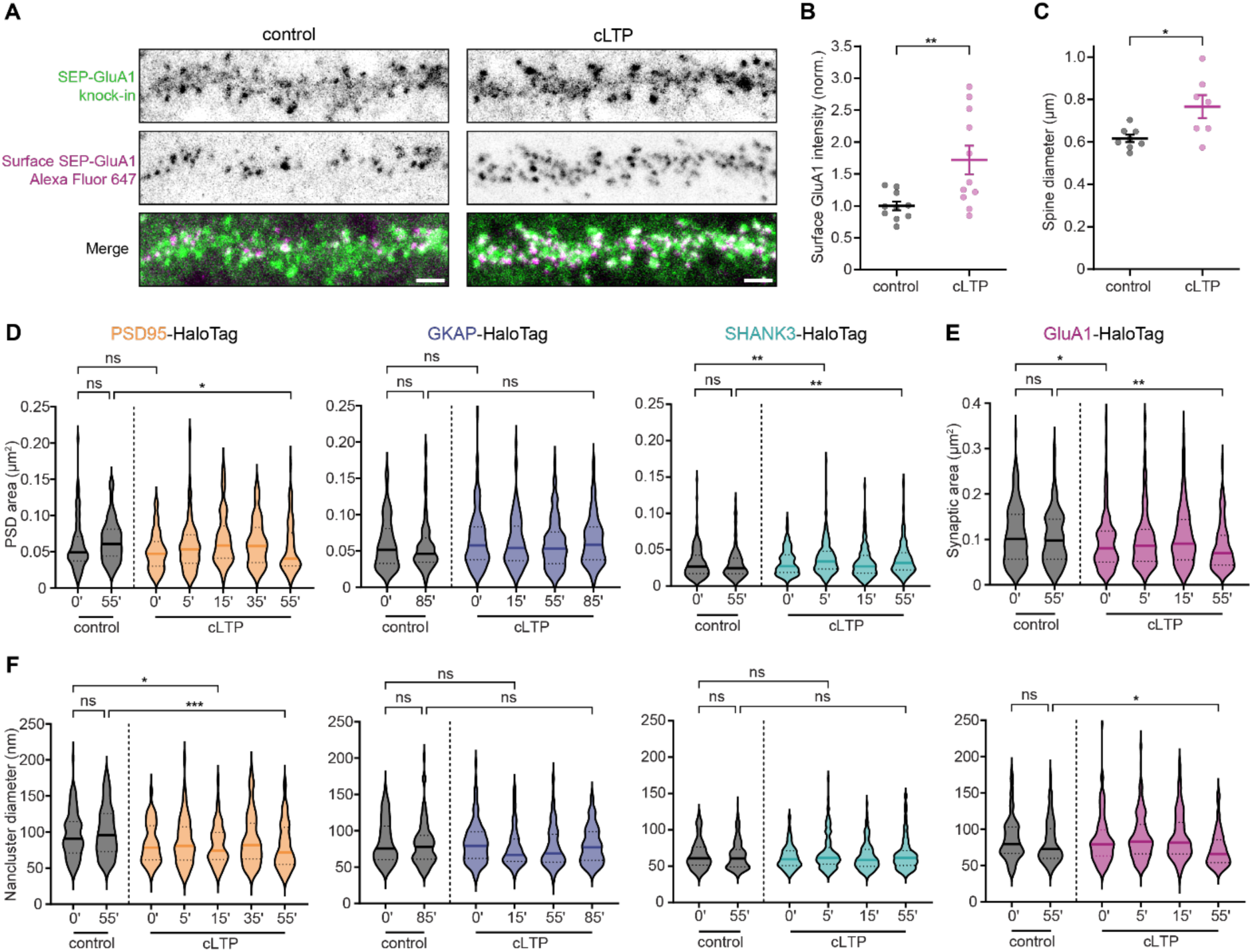
cLTP induces an increase in surface GluA1-containing AMPA receptors and spines size, but no robust increase in PSD area within 55 min. **(A)** Representative examples of hippocampal dendrites 55 min after cLTP induction. Shown are the SEP-GluA1 knock-in and surface SEP-GluA1 using confocal microscopy. Images are crops of maximum-intensity projections of acquired z stacks. Scale bar: 2 µm. **(B)** Quantification of surface SEP-GluA1 levels in control and cLTP conditions. Sample size: N=10-11 neurons. **(C)** Quantification of spine diameter in control and cLTP conditions. Sample size: N=8-7 neurons. **(D)** Violin plot of PSD area based on PSD95, GKAP and SHANK3 knock-ins over time after cLTP induction. DBSCAN was used to determine the PSD cluster in the dSTORM data. Sample size: N=80-208 synapses from 10-18 neurons. **(E)** Area of synaptic GluA1 based on GluA1-HaloTag over time after cLTP induction. DBSCAN was used to determine the synaptic cluster in the dSTORM data. Sample size: N=126-182 synapses from 9-13 neurons. **(F)** Violin plot of diameter of PSD95, GKAP, SHANK3 and GluA1 nanoclusters over time after cLTP induction. Sample sizes are the same as in D and E. Data in B and C are analysed using unpaired t-test. Data in D-F are analysed using Kruskal-Wallis test. Post-hoc comparisons are done using Dunn’s test. \**p* < 0.05, \*\**p* < 0.01 and \*\*\**p* < 0.001.

**Figure S3:**
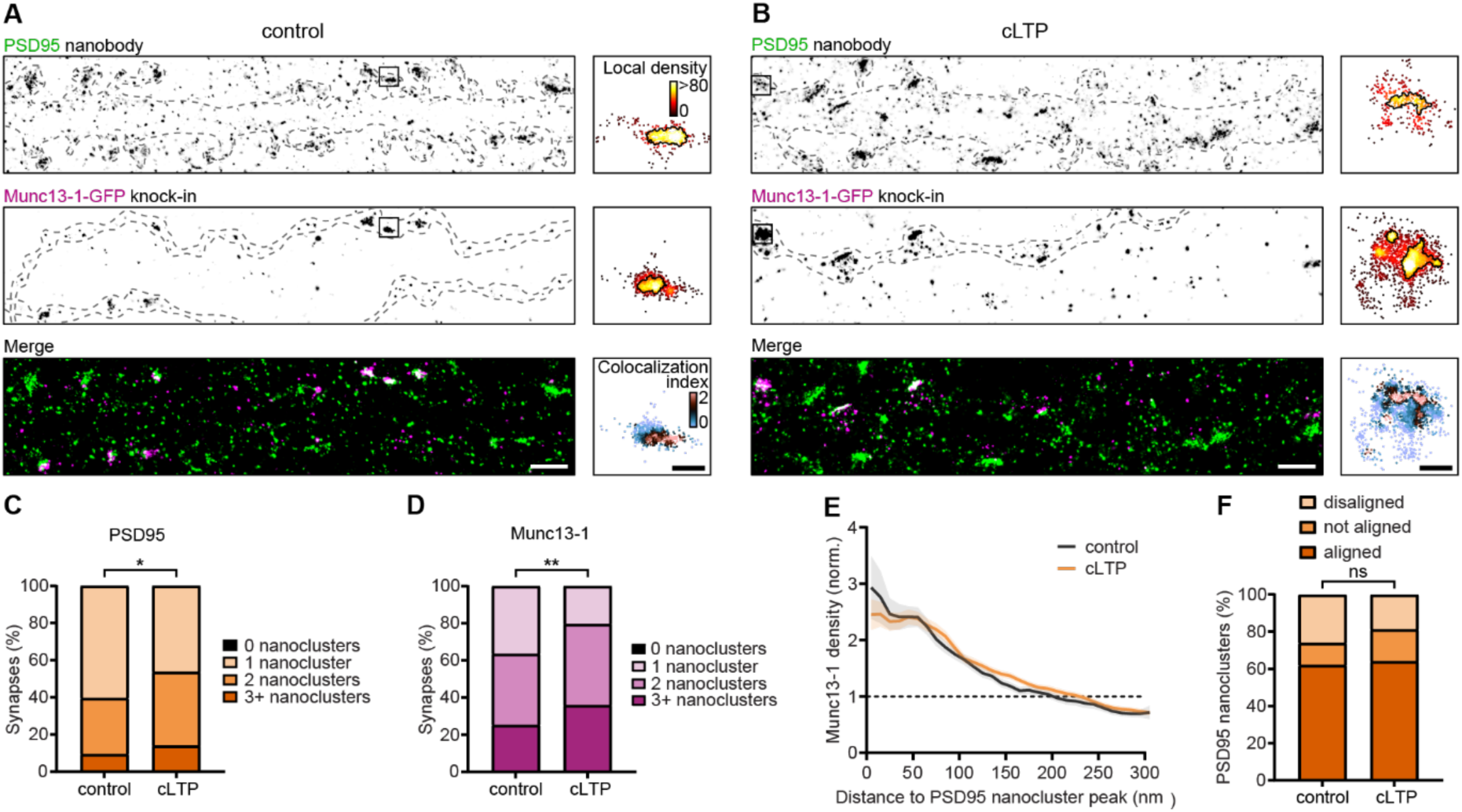
Newly formed Munc13-1 nanoclusters are aligned with PSD95 nanoclusters. **(A-B)** Representative examples of hippocampal dendrites and axons 55 min after control (A) or cLTP (B), marked by a PSD95 staining using a nanobody and endogenously labelled Munc13-1, respectively (scale bar: 2 µm). An example PSD and active zone after nanocluster analysis are given to the right of the dSTORM reconstruction (scale bar: 200 nm). The colocalization index of Willems et al. (2022)^1^ is used to visualize colocalization when PSD95 and Mun13-1 are overlayed. **(C)** The number of PSD95 nanoclusters 55 min after cLTP. Sample size: N=117-121 synapses from 21-22 neurons. **(D)** The number of Munc13-1 nanoclusters 55 min after cLTP. Sample sizes are the same as in C. **(E)** Munc13-1 localization density as a function of distance to the peak of PSD95 nanoclusters, normalized to randomized localisations. **(F)** Stacked bar graph of disaligned, not aligned and aligned PSD95 nanoclusters. Nanoclusters were considered aligned or disaligned if the enrichment of Munc13-1 localizations inside PSD95 nanoclusters was 1.96 standard deviations greater or smaller than randomized localizations, respectively. Kolmogorov-Smirnov test, p=0.684. Data in C and D are analysed using Mann-Whitney U test. \**p* < 0.05, \*\**p* < 0.01 and \*\*\**p* < 0.001.

**Figure S4:**
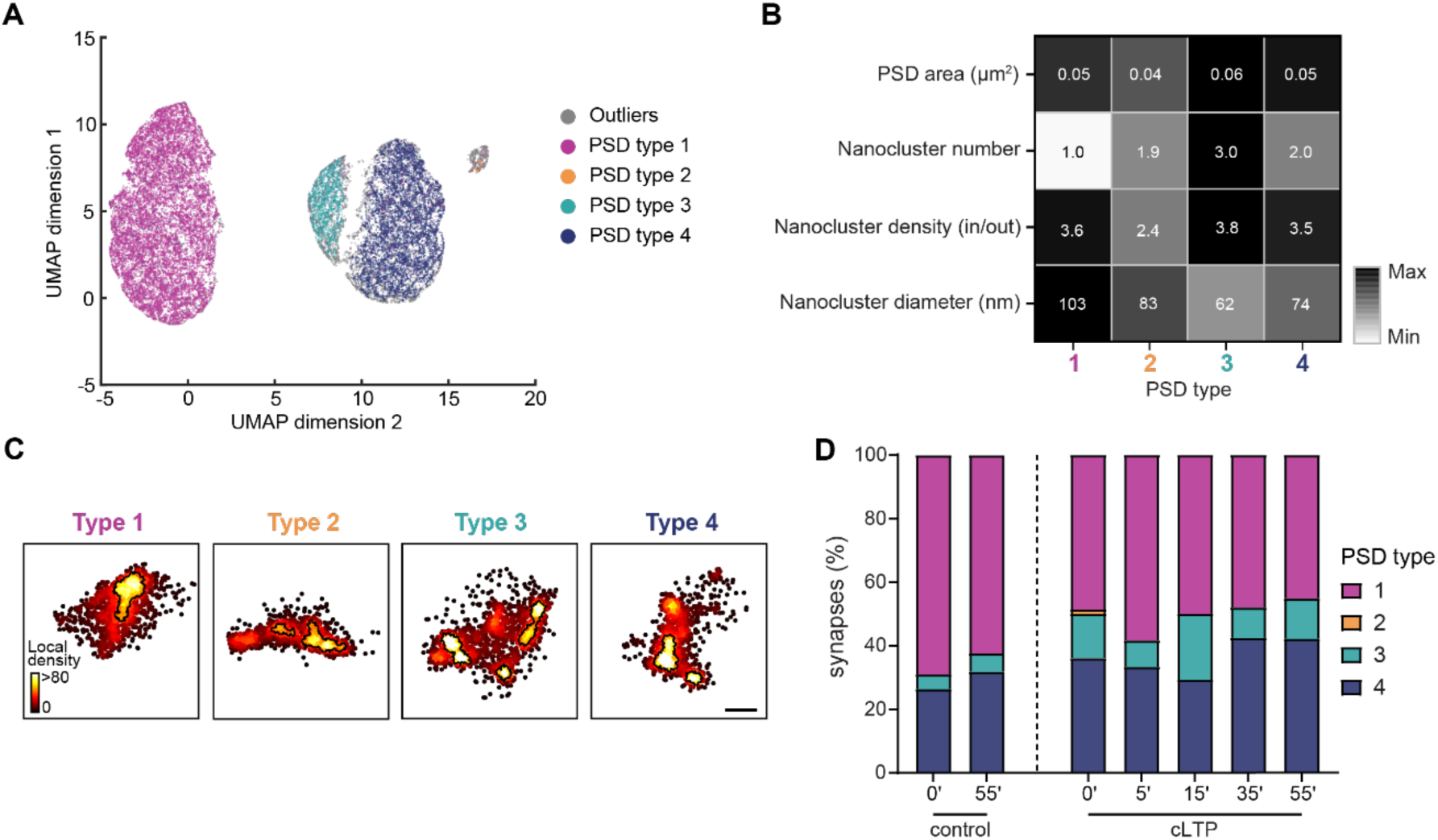
PSD nano-organization becomes more complex after cLTP induction. **(A)** UMAP representation of all PSDs analysed using HDBSCAN. 40,280 PSDs marked by PSD95-HaloTag were clustered based on their nano-organization (see Methods). **(B)** Heat map of normalized mean values of nano-organization of the identified PSD types. Mean values of PSD area (µm^2^), nanocluster number, nanocluster density (in/out) and nanocluster diameter (nm) are displayed for each PSD type in the corresponding square. **(C)** Representative examples of each PSD type visualized using dSTORM imaging of PSD95-HaloTag labelled with JF646. Scale bar: 200 nm. **(D)** Stacked histogram of the frequency of each PSD type after cLTP induction.

**Figure S5:**
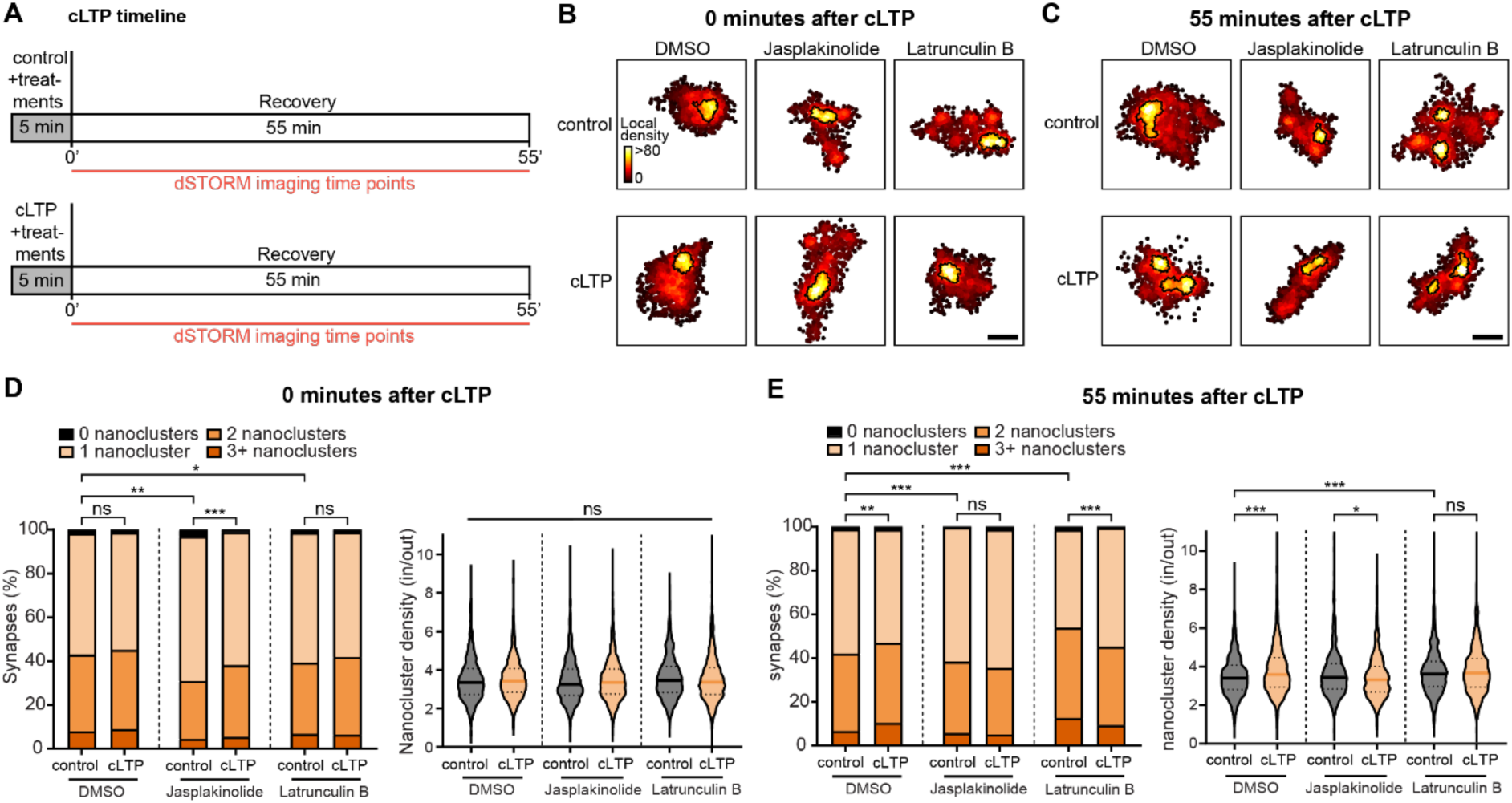
F-actin stabilization occludes cLTP-induced PSD reorganization, but actin depolymerization and recovery facilitate the formation of high-density nanoclusters. **(A)** Timeline of cLTP experiment. Dissociated hippocampal neurons (DIV 21) were treated with DMSO, 0.2 µM Jasplakinolide or 1 µM Latrunculin B during cLTP induction. Neurons were fixed either directly or 55 min after cLTP induction. **(B)** Representative examples of PSDs directly after cLTP induction, visualized using dSTORM imaging of PSD95-HaloTag labelled with JF646. Scale bar: 200 nm. **(C)** Representative examples of PSDs 55 min after cLTP induction. Scale bar: 200 nm. **(D)** Number of PSD95 nanoclusters per PSD and nanocluster density 0 min after cLTP induction and drug treatments. Left: Stacked histogram of the number of PSD95 nanoclusters per PSD. Sample size: N= 830-2066synapses from 7-15 neurons. **(E)** Number of PSD95 nanoclusters per PSD and nanocluster density 55 min after cLTP induction and drug treatments. Left: Stacked histogram of the number of PSD95 nanoclusters per PSD. Sample sizes were the same as in D. Data are analysed using Kruskal-Wallis test. Post-hoc comparisons are done using Dunn’s test. \**p* < 0.05, \*\**p* < 0.01 and \*\*\**p* < 0.001.

**Figure S6:**
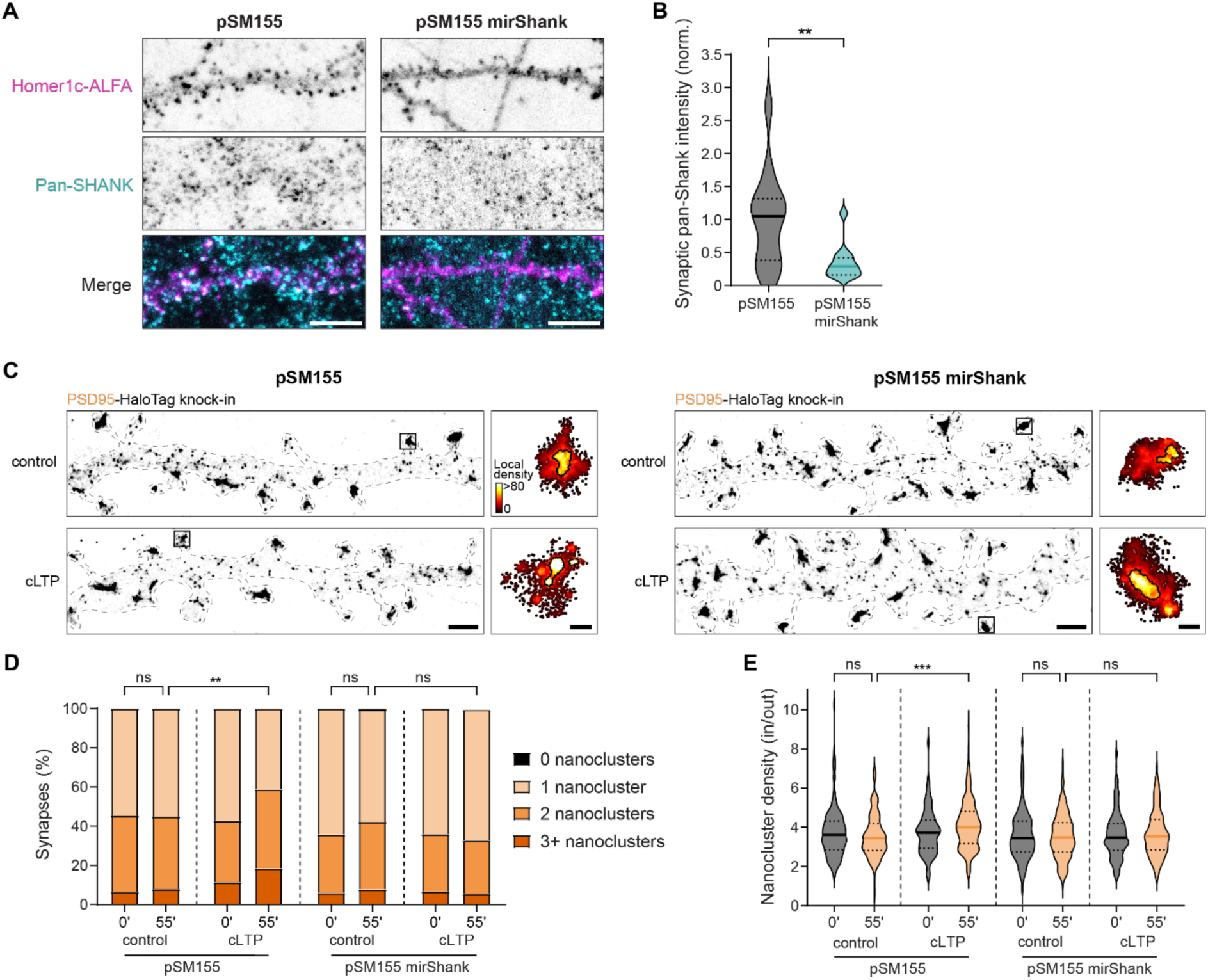
Knock-down of SHANK proteins occludes cLTP-induced PSD reorganization. **(A)** Dissociated rat hippocampal neurons were transfected at DIV 4 with the PSD95-HaloTag knock-in and either empty pSM155 mCherry or pSM155 mirShank::mCherry. Shown are representative examples of pan-SHANK signal in dendrites (DIV 21) in control and mirShank conditions. Homer1c-ALFA is visualized with Atto488 and is used as a synaptic marker. Scale bar: 5 µm. **(B)** Quantification of pan-SHANK levels in synapses marked by Homer1c in control and mirShank conditions. Mann-Whitney test, *U*=34, *P*=0.009. Sample size: N=12-14 neurons. **(C)** Representative dendritic stretches with highlighted PSDs (after nanocluster analysis) of reconstructed dSTORM images of PSD95-HaloTag labelled with JF646, 55 min after cLTP induction. Scale bar left: 1 µm. Scale bar right: 200 nm. **(D)** Stacked histogram of PSD95 nanocluster number per PSD 55 min after cLTP induction. Sample size: N=130-182 synapses from 10-14 neurons. **(E)** Violin plot showing the density of PSD95 nanoclusters after cLTP with and without SHANK knock-down. Kruskal-Wallis test, *H*=27.98, *P*<0.001. Dunn’s test, pSM155 55 min control versus pSM155 55 min cLTP: *P*<0.001. Sample sizes were the same as in D. Data in D and E are analysed using Kruskal-Wallis test. Post-hoc comparisons are done using Dunn’s test. \**p* < 0.05, \*\**p* < 0.01 and \*\*\**p* < 0.001.

**Figure S7:**
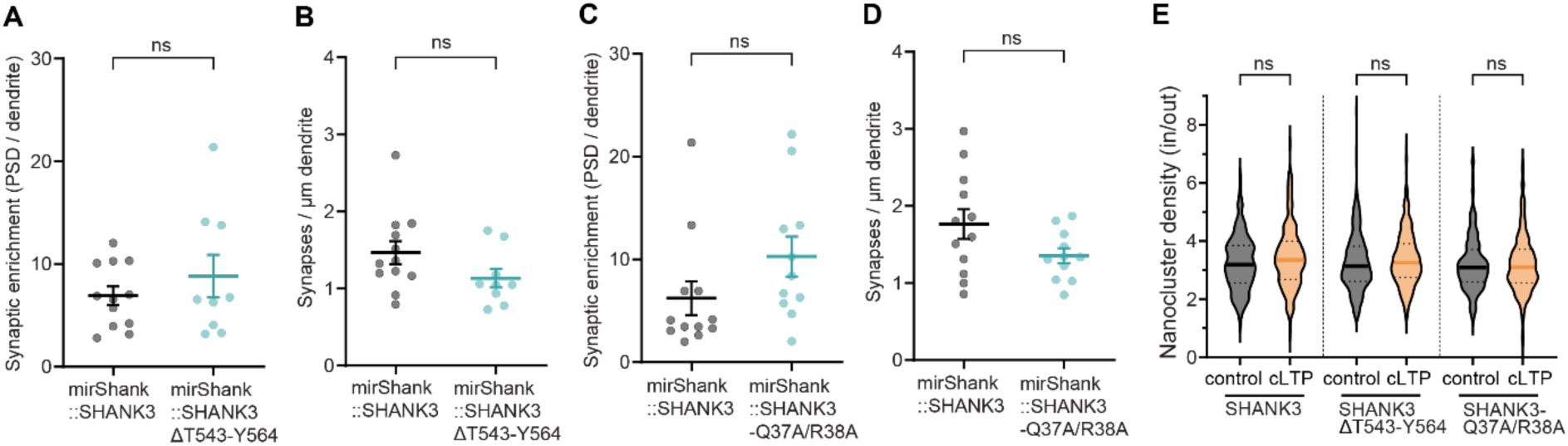
SHANK3ΔT543-Y564 and SHANK3-Q37A/R38A are enriched at the synapse and do not influence synapse number, PSD area, and nanocluster size. **(A-D)** Dissociated rat hippocampal neurons were transfected with Homer1c-ALFA and mirShank::mCherry-SHANK3, mirShank::mCherry-SHANK3ΔT543-Y564 or mirShank::mCherry-SHANK3-Q37A/R38A at DIV 4 and fixed at DIV 21. Homer1c-ALFA was visualized with Atto488 and used as a synapse marker. **(A)** Synaptic enrichment of SHANK3ΔT543-Y564 compared to SHANK3. Sample size: N=9-12 neurons. **(B)** Number of synapses marked by Homer1c per µm dendrite. Same sample size as in **A**. **(C)** Synaptic enrichment of SHANK3-Q37A/R38A compared to SHANK3. Sample size: N=10-12 neurons. **(D)** Number of synapses marked by Homer1c per µm dendrite. Sample size was the same as in C. **(E)** Dissociated rat hippocampal neurons were transfected at with the *PSD95*-HaloTag knock-in and mirShank::mCherry-SHANK3, mirShank::mCherry-SHANK3ΔT543-Y564, or mirShank::mCherry-SHANK3-Q37A/R38A. Shown is a violin plot of PSD area based on PSD95 55 min after cLTP induction. Sample size: N=209-304synapses from 11-16 neurons. Data in A-D are analysed using Unpaired t-test. Data in E are analysed using Kruskal-Wallis test. Post-hoc comparisons are done using Dunn’s test. \**p* < 0.05, \*\**p* < 0.01 and \*\*\**p* < 0.001.

**Table S1.**
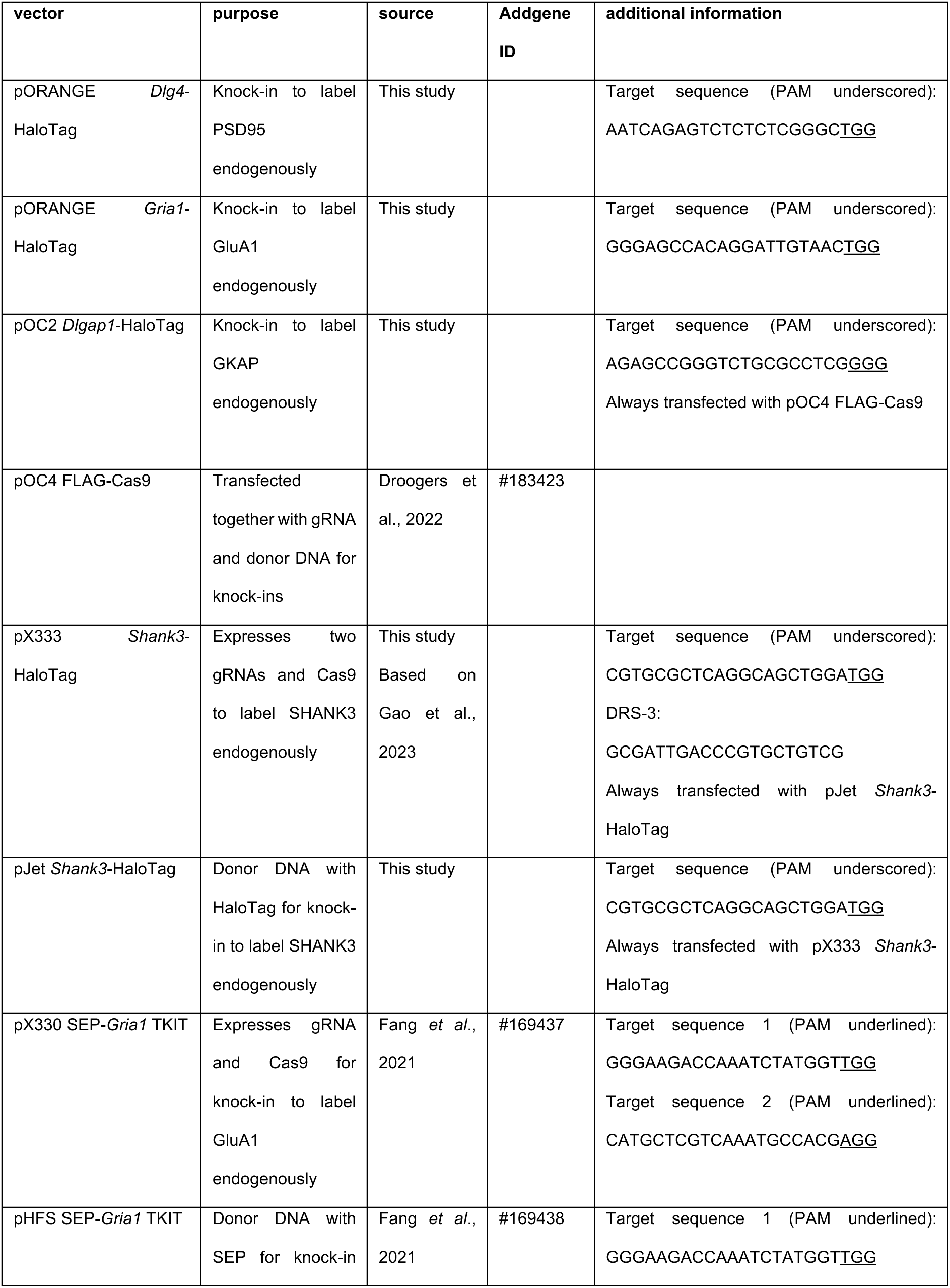
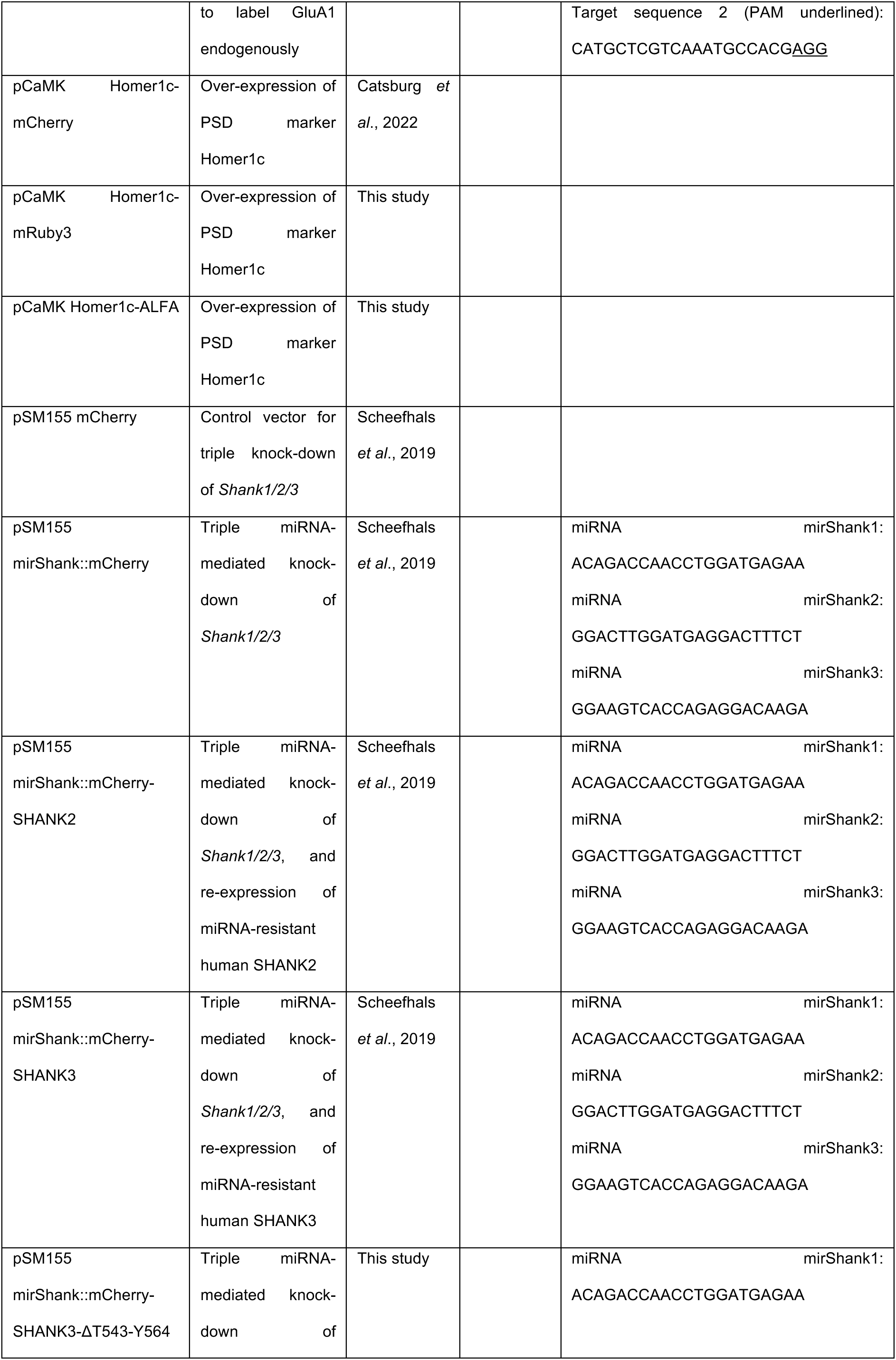
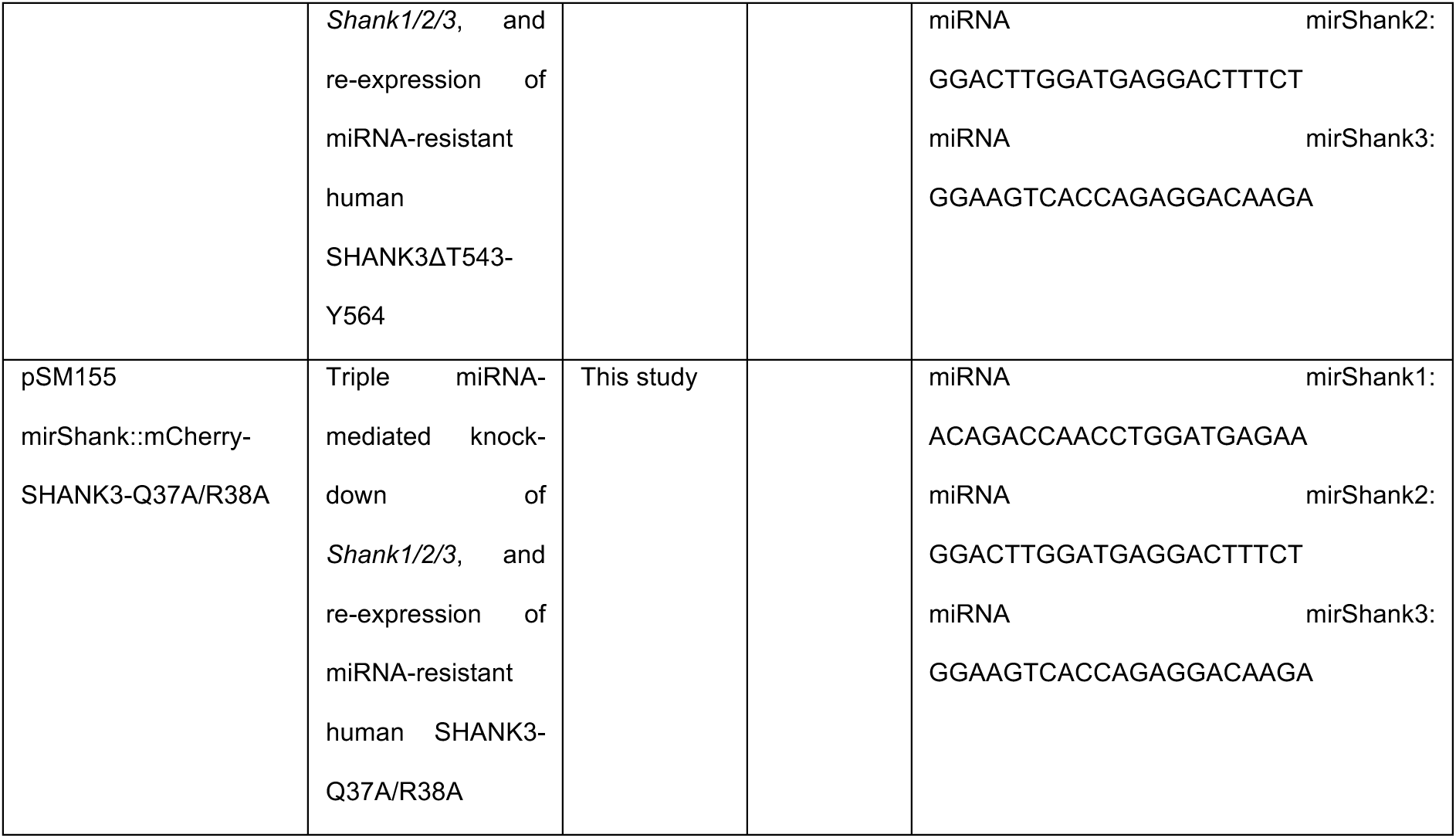
Vectors. All vectors that were used in this study are indicated with their name, purpose, source, Addgene ID, and additional information.

**Table S2.**
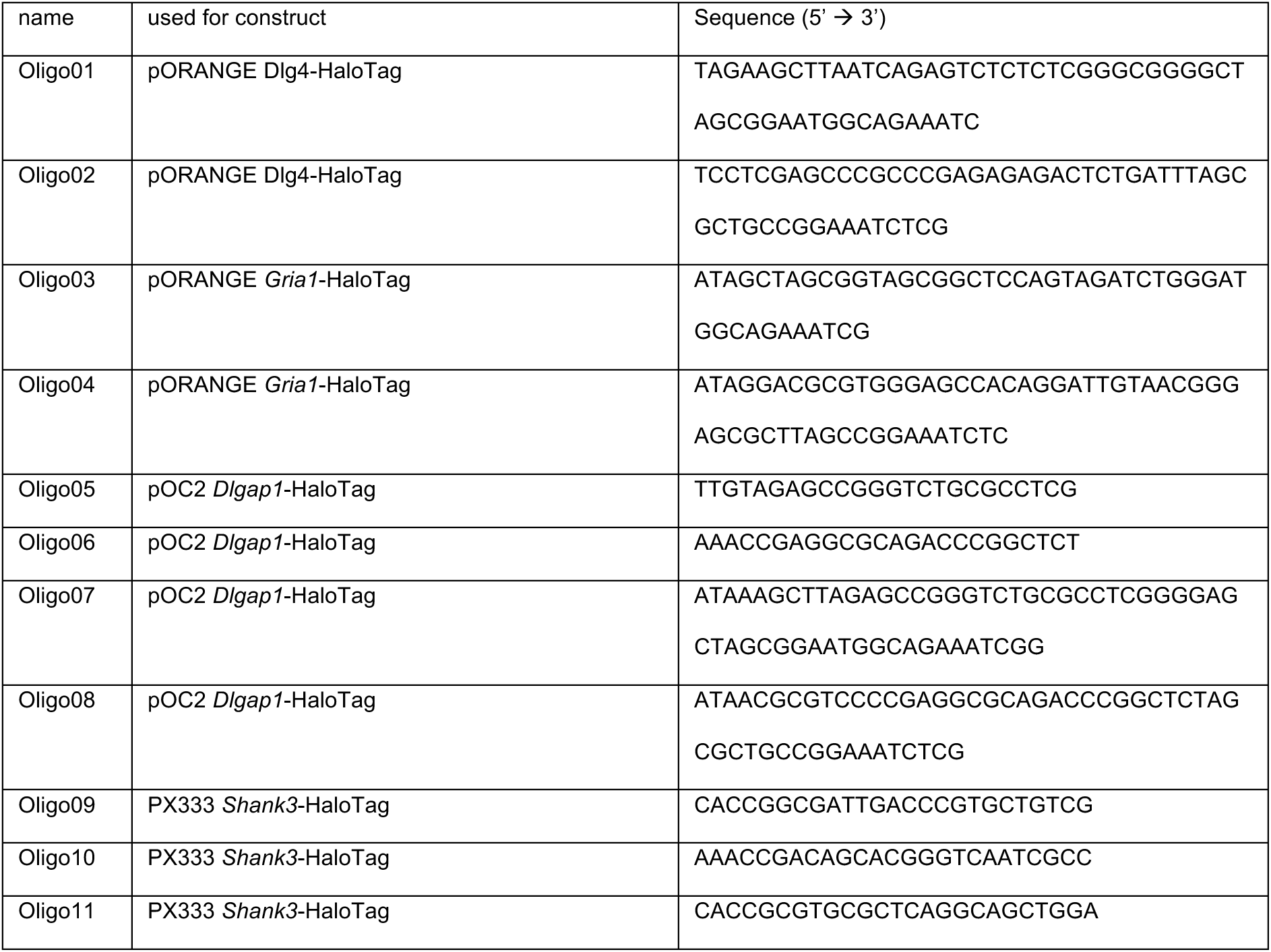
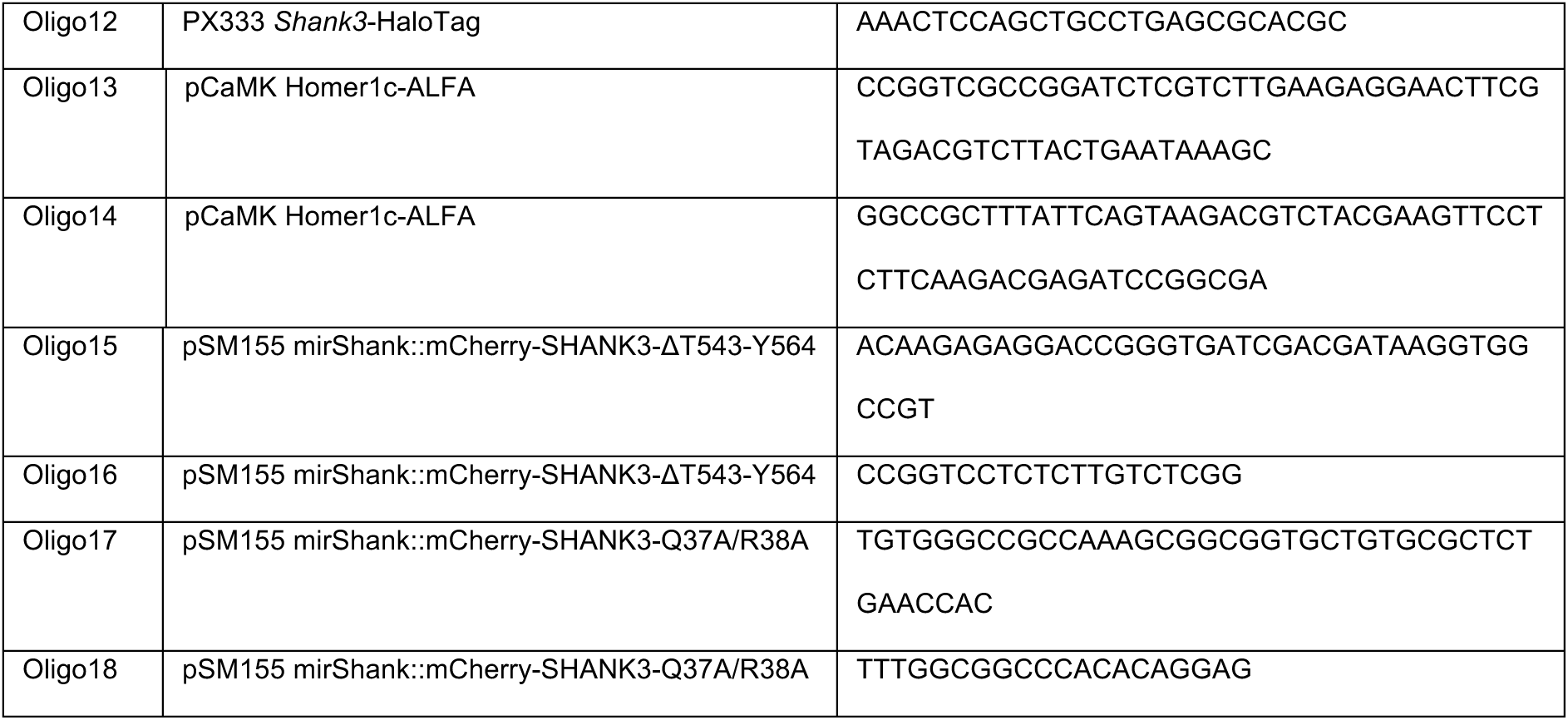
Oligonucleotides. All primers used to construct the vectors listed in Table1.

**Table S3.** Antibodies and reagents. All antibodies and reagents used are listed with their source, RRID or catalog number, and working concentration.

| **name** | **source** | **idenfifier** | **working concentration** |
|---|---|---|---|
| Anti-GFP mouse (IgG2a) | Thermo Fisher Scientific | AB_221568 | 1:2000 |
| Anti-pan-SHANK mouse (IgG1) | Antibodies Inc. | AB_10672418 | 1:500 |
| Anti-ALFA rabbit | NanoTag | AB_3075998 | 1:500 |
| Anti-ALFA Atto488 | NanoTag | AB_3075982 | 1:500 |
| Anti-mouse Alexa Fluor 647 | Invitrogen | AB_2535805 | 1:800 |
| Anti-mouse Alexa Fluor 488 | Invitrogen | AB_2534088 | 1:800 |
| Anti-rabbit Alexa Fluor 405 | Invitrogen | AB_2890548 | 1:1000 |
| JFX554 HaloTag® Ligand | Janelia Research Campus | Generous gift from Luke Lavis | 0.2 µM |
| JF646 HaloTag® Ligand | Promega | Cat. No. GA1120 | 0.2 µM |
| Bicuculline methiodide | Tocris | Cat. No. 2503 | 25 µM |
| DL-AP5 | Tocris | Cat. No. 0105 | 200 µM |
| KN-93 | Millipore | Cat. No. 422711-M | 5 µM |
| Jasplakinolide | Tocris | Cat. No. 2792 | 0.2 µM |
| Latrunculin B | Tocris | Cat. No. 3974 | 1 µM |
| Catalase | Sigma Aldrich | Cat. No. C40 | 40 µg/mL |
| Glucose oxidase | Sigma Aldrich | Cat. No. G2133 | 700 µg/mL |
| Cysteamine | Sigma Aldrich | Cat. No. 30070 | 5-35 mM |

**Table S4.** Reaction rates of AMPA receptor activation. Forward and backward rates of the 7 different AMPA receptor states of the MCell4 model of the synapse used in Figure 5. C0, C1, C2, C3, C4 and C5 are closed states. O is the open state of AMPA receptors.

| **reaction** | **forward rate (sec<sup>-1</sup>)** | **backward rate (sec<sup>-1</sup>)** |
|---|---|---|
| AMPAR C0 + glutamate ←→ AMPAR C1 | 4.59 * 10<sup>6</sup> | 4.26 * 10<sup>3</sup> |
| AMPAR C1 + glutamate ←→ AMPAR C2 | 2.84 * 10<sup>7</sup> | 3.26 * 10<sup>3</sup> |
| AMPAR C2 ←→ AMPAR O | 4.24 * 10<sup>3</sup> | 900 |
| AMPAR C1 ←→ AMPAR C3 | 2.89 * 10<sup>3</sup> | 39.2 |
| AMPAR C2 ←→ AMPAR C4 | 172 | 0.73 |
| AMPAR C3 + glutamate ←→ AMPAR C4 | 1.27 * 10<sup>6</sup> | 45.7 |
| AMPAR C4 ←→ AMPAR C5 | 16.8 | 190.4 |
| AMPAR C5 ←→ AMPAR O | 17.7 | 4 |

## Notes

### Competing Interest Statement

The authors have declared no competing interest.

